# Distinguished biological adaptation architecture aggravated population differentiation of Tibeto-Burman-speaking people inferred from 500 whole-genome data from 39 populations

**DOI:** 10.1101/2023.06.16.545243

**Authors:** Yuntao Sun, Mengge Wang, Qiuxia Sun, Yan Liu, Shuhan Duan, Zhiyong Wang, Yunyu Zhou, Jun Zhong, Yuguo Huang, Xinyu Huang, Xiangping Li, Haoran Su, Yan Cai, Xiucheng Jiang, Jing Chen, Jiangwei Yan, Shengjie Nie, Liping Hu, Junbao Yang, Renkuan Tang, Chuan-Chao Wang, Chao Liu, Xiaohui Deng, Guanglin He, Libing Yun

## Abstract

Tibeto-Burman (TB) people have tried to adapt to the hypoxic, cold and high-UV high-altitude Tibetan Plateau and complex disease exposure in the lowland wet and hot rainforest since the late Paleolithic period. However, the full landscape of genetic history and biological adaptation of geographically diverse TB people and their interaction mechanism remained unknown. We generated a whole-genome-based meta-database of 500 individuals from 39 TB populations from East Asia and Southeast Asia and presented a comprehensive landscape of genetic diversity, admixture history and differentiated adaptative features of geographically different TB people. We identified geography/language-related genetic differentiation among Tibetan Plateau, Tibetan-Yi-Corridor (TYC) and Southeast Asian TB people, consistent with their differentiated admixture process with incoming or indigenous ancestral source populations. A robust genetic connection between TYC people and ancient YR people supported the Northern origin hypothesis of TB people. We reported population substructure-related differentiated biological adaptative signatures between highland Tibetans and lowland TB people and between geographically different Lolo speakers. Highland adaptative *EPAS1* and *EGLN* variants riched in Tibetans but lacked in TYC people whose adaptation is associated with the physical features and skin pigmentation (*EDAR* and *SLC24A5*), hepatic alcohol metabolism (*ALDH9A1*), regulation of cell-cell adhesion of muscle cells (*CTNNA3)* and immune/fat metabolism-related adaptative signature. TB-related genomic resources provided new insights into the genetic basis of phenotype differences and better reference for the anthropologically-informed sampling design in biomedical and genomic cohort research.

## INTRODUCTION

Tibeto-Burman (TB)-speaking people possess high ethnolinguistic diversity and live in areas with complex terrain, ranging from Tibetic-speaking Tibetan and Sherpa people from the Tibetan Plateau (TP) to Burmese-speaking people from lowland South China and Southeast Asia ^1^. The Bodic speakers are widely distributed in highland East Asia, and Lolo-Na-Qiangic-speaking populations, mainly including Yi, Qiang, Bai, Lahu and Kham Tibetans, reside in the Tibetan-Yi corridor (TYC) and the Yunnan-Guizhou Plateau (YGP) in Southwest China. The Burmese-speaking ethnic groups, including Achang and others, live in mainland Southeast Asia (MSEA). Previous linguistic, archaeological and genetic evidence have evidenced that TB/Sinitic-speaking people have a common origin and drawn a glorious history of Chinese civilization ^2, 3^. Differentiated local biological adaptation and geographical/cultural barriers, including genetic isolation caused by the river or mountains, have permanently influenced the genetic profiles of ethnic groups in these areas ^4, 5^. Further extensive population migration and complex admixture with other East Asians, including Altaic, Tai-Kadai (TK), Hmong-Mien (HM), Austronesian (AN) and Austroasiatic (AA) speakers, further complexed the general patterns of TB people. A previous study revealed that geographically different but ethnically similar TB people shared differentiated genetic backgrounds, such as genetic differentiation between Ü-Tsang Tibetans in the core-Tibet region, Sherpa people in the southern slope of the Himalayas, and Ando Tibetans in Gansu-Qinghai area ^6^. However, the overall patterns of genetic similarities and differentiations of geographically different TB people and their detailed interaction with other ancient and modern East Asians remained unknown.

The genetic origin of modern TB people attracted the attention of scientists from different research areas. Archaeological evidence suggested that Yangshao and Qijia cultures from Northern China’s Yellow River (YR) millet agriculture were related to the Proto-TB people ^7–9^. Early linguistic evidence has provided clues for the origin from North India, western Sichuan and North China related to the North India Hypothesis, TYC Hypothesis and Northern-China origin hypothesis. Zhang and other linguistic scientists recently reconstructed linguistic phylogeny relationships of the TB language family, supporting the third hypothesis associated with the ancient Neolithic Yangshao and Longshan cultures from North China ^3, 10, 11^. Previous genetic research has provided substantial evidence for the admixture trajectory of highland TB people, mainly for Tibetan and Sherpa ^12, 13^. Lu et al. illuminated two-layer ancestral components of modern Tibetans ^13^, and Zhang et al. comprehensively characterized the genetic differentiation between highland Tibetan and Sherpa people ^12^. Genetic differences among geographically different highland TB people and their solid genetic connection with lowland northern East Asians were also confirmed via the genome-wide array data ^6, 14^ and forensic-related Insertion/Deletion ^15^, autosomal single nucleotide polymorphisms and short tandem repeat ^16, 17^. Early large-scale genetic evidence from paternal and mitochondrial variations confirmed that the archaeologically and linguistically supported population migration and admixture events contributed to forming the uniparental gene pool of TB people ^18, 19^. Previous studies focusing on haplogroups D-M174 and O-M117 demonstrated that TB minorities result from admixture between Neolithic northern YR immigrants and Southern indigenous inhabitants ^20, 21^. Zhao et al. identified Paleolithic colonization of the TP and Neolithic expansion from Northern China based on a rare haplogroup M16 and other uniparental lineages ^22^.

The genetic history of TB in lowland regions from South China and Southeast Asia also provided new insights into the overall formation patterns of TB people. Kutanan et al. reported the genetic differentiation between TB people and other populations from Southeast Asia based on genome-wide SNP data and whole-Y sequencing variations ^23–25^. Ancient DNA has illuminated multiple gene flow events from southern Chinese rice farmers to ManBac-related ancient Southeast Asians and millet farmers to Oakaie-related ancient Myanmar people, providing a direct link between northern and southern TB people ^26^. Exome sequence evidence reported by Yang et al. demonstrated that language-related population stratifications in Yunnan were associated with the ancient origin of Baipu, Baiyue and Proto-TB people ^27^. However, the TYC was a critical geographical corridor linking highland TB and lowland Southeast Asia; only two genetic studies focused on the genetic structure were performed based on a small sample size of targeted ethnic groups ^5, 28^. Yao et al. found a mixed pattern of northern TYC TB speakers, who derived their ancestry mainly from Tibetan and Han ^28^. Zhang et al. explored the differentiated genetic landscape of geographically different TYC people based on low-coverage (∼5X) whole-genome sequencing (WGS) data ^5^. However, the complexity of the genetic diversity of TYC people inferred from the forensic-related markers showed the missing diversity and gap in the evolutionary history of linguistically diverse TYC populations at the whole-genome level ^29, 30^. Generally, genomic resources from TP and surrounding regions highlighted the unique genetic structure and distinctions among geographically separated TB populations ^31–33^. We also noticed that the genomic footprints of the evolution of TB people were portrayed incomprehensively, such as the entire landscape of patterns of genetic diversity caused by sampling bias, small sample size, and coverage of single areas. Another critical limitation of previous genetic studies was the lack of comprehensive population comparisons of publicly available TB people and an entire genetic landscape of geographically different TB people.

Understanding human genomic diversity and biological adaptative processes is fundamental to discovering the association between genetic variations, complex physiological traits and genetic disease susceptibility ^34^. Recently China has proposed multiple Human Genome Cohort Projects such as NyuWa genomic resource ^35^, 10K Chinese People Genomic Diversity (10K_CPGDP) ^36^, Westlake BioBank for Chinese (WBBC) ^37^, 100K Genome sequencing for Rare Disease (GRSD^100K^) and China Metabolic Analytics Project (ChinaMAP) ^38^. Nevertheless, the genetic diversity of TB-speaking ethnic populations remains not characterized at a fine scale ^39–42^. As mentioned, TB people had the feature of a large population, wide geographical distribution and significant genetic differentiation. Current studies still had two significant limitations: (1) restrictions in representative sampling ethnic groups, i.e., most reported data preferred Tibetans from the TP ^43–46^, leading to inaccurate surveys of the fine-scale genetic structure to onefold ethnic ^47^. (2) the lack of comprehensive geographical coverage of different ethnical samples, i.e., the published studies adopted regional data without combining geographical distinction information to refine TB genomes’ footprints ^5, 24, 27^. To overcome these limitations, we constructed entire-regional meta-databases of TB populations and systematically characterized their trait-related adaptive variants and genes. We combined newly generated data with previously published data from all TB-speaking people, including Ü-Tsang Tibetans from the core-Tibet region in the TP; Ando Tibetans residing in Gangcha, Gannan and Xunhua from the Gansu-Qinghai region; Kham Tibetans from Xinglong, Yajiang and Yunnan and other TYC groups; Pumi, Naxi, Qiang, Bai, Lahu, Guizhou Yi (GZY) living in Southwest China, Tujia in Eastern lowland as well as Burmese-speaking ethnic groups in MSEA ^3, 6, 24, 25^ (**Figure S1 and Table S1**). With comprehensive population genetic analyses based on shared alleles and phased haplotypes of 500 individuals from 39 populations, we aimed to portray the panorama of population structure, demographic history and local adaption of TB populations. We identified differentiated genetic structure and adaptative features of geographically different TB people, which were influenced by the geographical and cultural barriers and evolutionary forces related to population admixture and selection under unique environments. We found apparent genetic substructures and different selection signals between ethnically close TB speakers (SCY and GZY living in different areas) and between ethnically distinct TB speakers (Tibetans in the TP and Yi people in the middle-altitude region). Our findings suggested that demographic events, including complex population admixture and differentiated adaptative features, combined with cultural and geographical barriers, such as ethnic customs of marriage and belief and historical events of war and policy, contributed to TB’s genetic differentiation and the different patterns of genetic diversity of ecologically different TB people.

## RESULTS

### General pattern of genetic affinity and differentiation of modern East Asians

We presented one merged meta-database, including publicly available and newly-generated modern East Asians and worldwide reference populations ^3, 6, 24, 25^, to explore the genetic background of TB-speaking populations. The clustering pattern was first inferred from the global principal component analysis (PCA), including Human Genetic Diversity Project (HGDP), Oceanian genomic resources and other populations ^48–50^. We found that the first two principal components (PC1 and PC2) revealed the continental-scale genetic differentiation of the worldwide populations, and TB ethnic groups overlapped with East Asians, showing a closer relationship with East Asians compared with non-East Asian groups (**Figure S2**). As anticipated from the PCA results, the estimates of pairwise Fst matrixes further showed the smallest differentiation between TB and East Asians (Fst ≤ 0.0664) (**Figure S3 and Table S2**). We found that geographically different TB people had less degree of differentiation (0.0007-0.0368) compared to other linguistically different ethnic groups, such as HM ethnic groups (0.0025-0.0559) and Altaic ethnic populations (0.0041-0.0560), which supported the relatively strong genetic affinity within TB people. We also identified the most considerable genetic distinction between Tibetan and TB people in Southeast Asia, larger than that between Tibetan and Han Chinese. We next constructed the Neighbor-Joining (NJ) tree based on the Fst matrix (**Figure S4**), which portrayed that the TB people clustered with East Asian groups but were divided into different subgroups by linguistically different Sinitic/HM/TK populations. Our results demonstrated that geographically separated TB groups showed different phylogenetic relationships with one another.

We herein assessed whether the TB people were separated into geography/language-related population stratifications. After removing other worldwide reference populations, we retained the TK-speaking populations as the representative of East Asians. The haplotype-based fineSTRUCTURE analyses (**Figure 1A**) showed the fine-scale genetic structure of 625 individuals. Three major genetic clusters were confirmed, one including Tibetic-speaking ethnic groups (Tibetan, Qiang and Pumi), another including Burmese-related and TK-speaking speakers, and the other including Sichuan Yi (SCY), GZY, Lahu and Tujia populations. ADMIXTURE analysis under the best-fitted model (K = 3) showed different patterns of the ancestral composition of the aforementioned clustered-based TB populations. Significantly, Burmese-speaking ethnic groups, Yi and Sinitic-speaking people derived more of their ancestry from southern East Asians related to TK-speaking people than Tibetic-speaking populations. The patterns of geography-related genetic substructure among TB groups were consistent with fineSTRUCTURE-based phylogeny.

**Figure 1.**
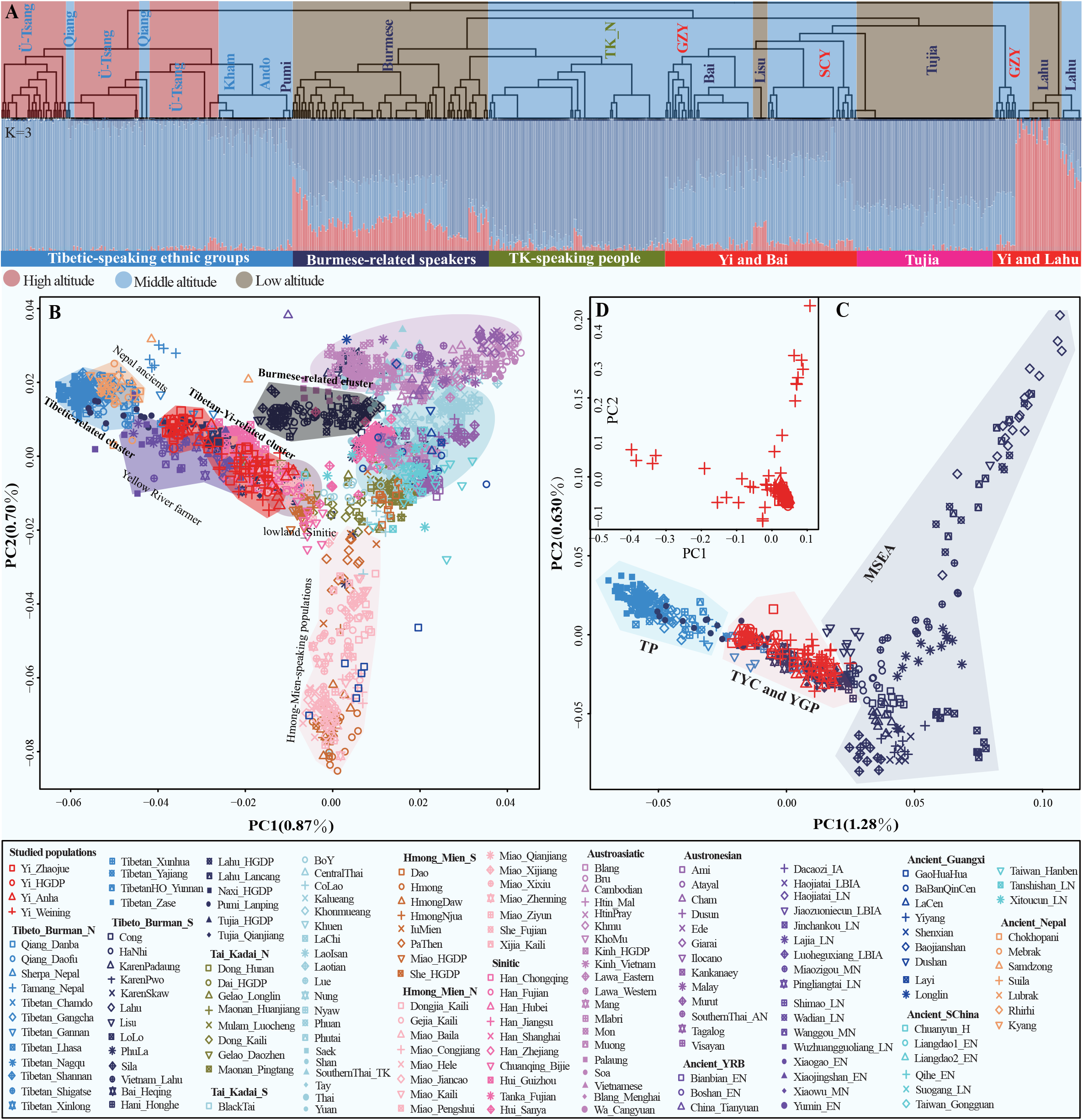
The genetic profile of geographically different TB populations. (**A**) The fineSTRUCTURE generated based on phased haplotypes of 625 individuals from 39 TB populations and TK people. Dendrogram shows the topological pattern and clustering among TB populations at high (red) - middle (blue) - low (brown) altitudes. ADMIXTURE shows the proportion of different ancestral components among different language-speaking populations. It illustrates the genetic substructure among TB populations. **(B-D)** Principal component analysis (PCA) at different scales demonstrates the clustering between TB populations and surrounding modern populations. Projections of Ancient_YRB, Ancient_South China, Ancient_Guangxi and Ancient_Nepal people to modern populations are presented, showing initial qualitative affinity. (**B**) PCA of modern and ancient people when restricted to East Asians. (**C**) PCA of TB populations at fine-scale. (**D**) PCA of four Yi population.

We further investigated the genetic affinity among TB populations via PCA at various scales. We observed that TB populations were distributed along the PC1 and formed three clusters in the East Asian context (**Figure 1B**). The Tibetan-related cluster included Ü-Tsang Tibetans, Ando Tibetans and Pumi populations. Populations from TYC and YGP, including SCY, GZY, Bai, Qiang and Naxi, were clustered into the Tibetan-Yi-related branch, while Karen, Lisu and Lolo from MSEA formed the Burmese-related branch. We further observed that populations from the same geographical area formed specific isolated clusters at a fine scale of PCA, including only TB populations (**Figure 1C**). The Tibetan subgroups (Pumi and Naxi), Ando Tibetans from Xunhua and Gannan, and Yajiang Kham Tibetans overlapped with Qiang, Bai and Yi populations distributed in TYC and YGP. In contrast, Burmese-speaking populations formed a cline alone, suggesting that TB populations from China were closer to each other than those from MSEA. These observed patterns were also supported by the results of Fst, outgroup-*f*_3_ and IBD (**Figures S5-6**). Next, we reconstructed the topology of TB populations based on the converted *f_3_*-related matrix (1-outgroup-*f*_3_). The phylogenetical relationships among TB populations were in line with different geographical patterns (**Figure S7**). The Sherpas and Tamang clustered with high-altitude Ü-Tsang Tibetans; TYC and YGP populations clustered together and formed the middle-altitude-related clade; Low-altitude populations (Burmese-speaking people and Tujia) clustered with each other. Our results identified that geographically different TB populations significantly differentiated from each other, and groups from the same geographical area harbored a closer genetic affinity, suggesting language/geography-associated population stratification existed in geographically different TB people.

### The influence of admixture on the patterns of genetic diversity of TB people

We conducted model-based ADMIXTURE to explore the basic patterns of admixture sources and proportions (**Figure 2A**). The well-fitted admixture model (K=8) in the context of East Asia revealed that TB groups had multiple ancestral compositions. The northern ancestral components appeared in TB people represented by Ü-Tsang Tibetans (red) and Lahu (blue). The southern source was associated with TK speakers represented by BoY (pink) and HM speakers represented by Hmong (light green). Northern TB derived primary ancestry from ancestors related to ancient TP people, and southern TB possessed more ancestry related to HM/TK-speaking people. It is worth noting that the proportions of ancestral components varied among geographically different TB populations. There was decreasing tendency of Ü-Tsang Tibetan-related ancestral component from high-altitude (ethnic groups from TP, TYC and YGP) to low-altitude populations (Tujia and Burmese speakers). This estimated result was confirmed by TB-based ADMIXTURE modeling (**Figure S8**).

**Figure 2.**
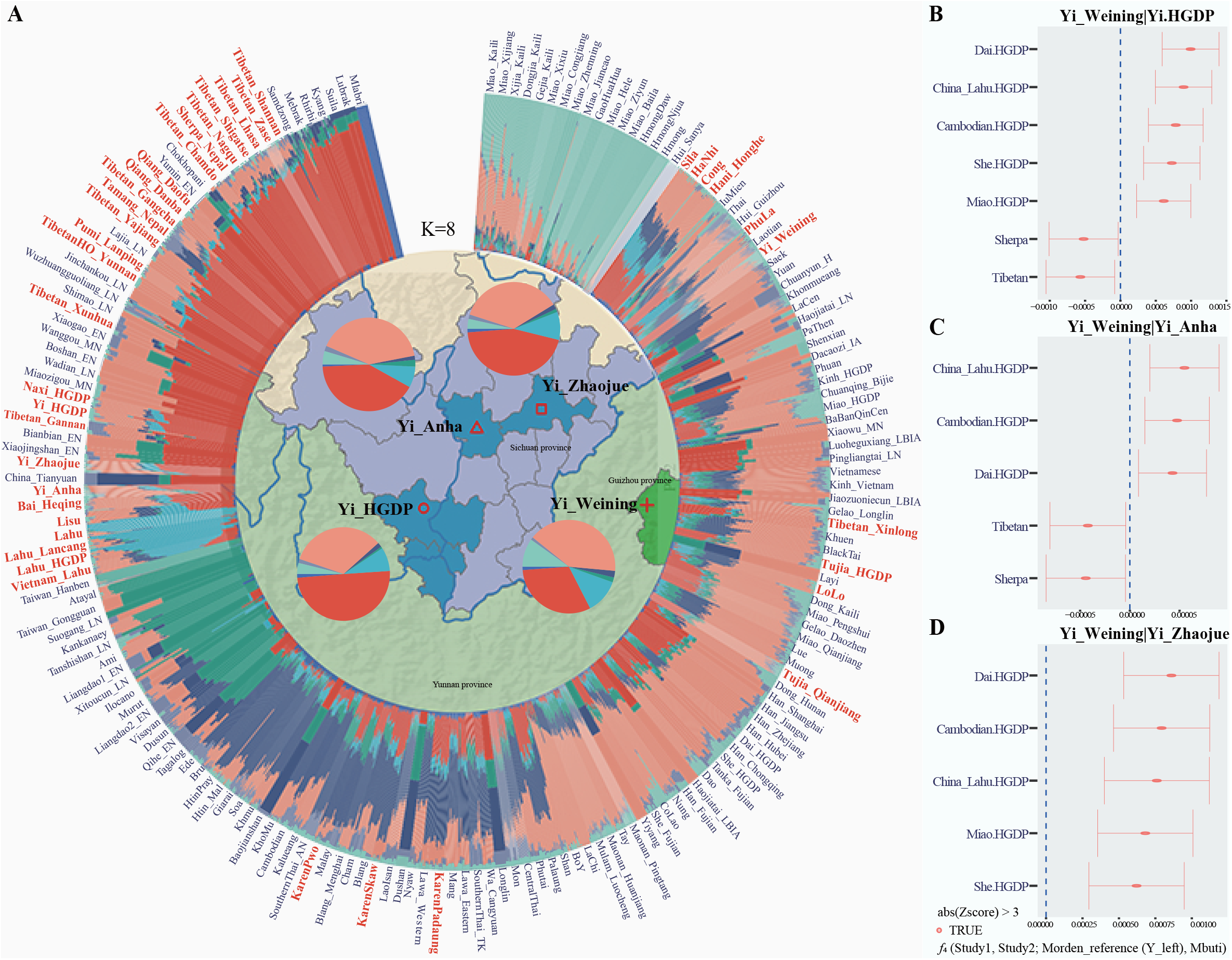
Ancestral composition and gene flow of the TB populations compared with the reference populations at the East Asian scale. **(A)** The outer circle is the result of ADMIXTURE chosen based on the minimum cross-validation (CV) error of eight ancestral populations (K=8), which shows the red composition of the northern Tibeto-Burman (TB) language group represented by Tibetans from the Tibetan Plateau (Ü-Tsang) and the Ancient Nepalese populations (aMMD), the blue composition of southern TB language group represented by Lahu, the light green composition of Hmong-Mien (HM) language family represented by Hmong, pink composition of Tai-Kadai (TK) language family represented by BoY, and the compositions of other language ethnics. The inner map shows the location of newly sequenced Yi populations and the ADMIXTURE results in the form of a pie chart. (B-D) Genetic distinctness between Yi from different areas and modern populations was explored based on *f_4_* tests, *f_4_*(GZY, SCY; Morden reference populations (Y_left), Mbuti), showing an approximate different geographical pattern of interaction between Yi and modern populations. That is, showing affinity between SCY (Yi_Zhaojue, Yi_Anha, Yi_HGDP) and surrounding TB populations, as well as the affinity between GZY (Yi_Weining) and surrounding ethnic groups such as HM and TK. As shown in the legend, absolute values of Z > 3 are indicated by TRUE, presenting differences between GZY and SCY. As shown in the legend, absolute values of Z > 3 are indicated by TRUE, presenting differences between GZY and SCY. The left side of the dotted X-axis represents negative values of f4, suggesting that SCY has gene flow events with modern reference populations compared to GZY. Similarly, the right side of the dotted X-axis represents positive values of f4, suggesting that GZY has gene flow events with modern reference populations compared to SCY.

We explored the potentially differentiated admixture that contributed to the observed genetic differentiation. We calculated symmetric *f*_4_-statistics in the form of *f*_4_(TB1, TB2; reference populations, Mbuti) and the first and second studied TB populations were considered to form a clade when |Z|<3. Positive Z-scores demonstrated that the reference populations shared more alleles with the first-studied populations than the second-studied populations and vice versa. We confirmed the differentiated shared ancestral components inferred from ADMIXTURE models that northern TB possessed more ancestry related to ancient northern East Asians and southern TB people shared more ancestry related to ancient southern East Asians. Focusing on our first-reported Yi people, GZY was genetically differentiated from SCY (Yi_Zhaojue, Yi_Anha and Yi_HGDP) in ADMIXTURE (**Figure 2A**). GZY received additional gene flow from southern Chinese and Southeast Asians (Dai, Lahu, Combodian, Miao and She) when compared to SCY. Similarly, SCY possessed additional gene flow events from highlanders (Tibetan and Sherpa) relative to GZY (**Figure 2B**). Substantial gene flow events among different source populations were also observed using a Treemix-based phylogenetic tree (**Figure S9A∼B**). ALDER-based two-way admixture results supported this admixture pattern with large-scale admixture time ranges with different source pairs (**Table S3**). Generally, the results above showed the different admixture histories of Yi populations. Geographically different Yi populations were affected by frequent admixture events from surrounding linguistically different populations, which contributed to their genetic differentiation.

### Genetic substructure and demographic analysis of ethnically similar groups

We found genetic differentiation between northern, central and southern geographically different TB people. We next focused on the fine-scale genetic substructure of underrepresented geographically diverse Yi populations. We observed a SCY-related cluster and two GZY-related clines (**Figure 1D**). When the ADMIXTURE was restricted to TB individuals, we observed intricate population substructures as ancestral sources (K)increased. Although the unique ancestral component of GZY was not found, their ancestral composition proportions were significantly different from SCY (**Figure S10**). To explore the fine-scale genetic structure of TB populations comprehensively, we phased 348 unrelated TB individuals and used fineSTRUCTURE to examine further the genetic affinity and differences among TB populations (**Figure 3A**). The reconstructed dendrogram split these populations into three major branches, color-marked as red, blue and purple, respectively. Three TYC populations (SCY, Naxi, and Pumi) were located at the same branch with highland Ü-Tsang Tibetan, GZY was situated at the same branch with other YGP populations, including Bai, Lahu and Hani, and Tujia individuals clustered together. It demonstrated the high genetic diversities of TB speakers from different geographical regions that contributed to the different genetic backgrounds. Another noteworthy observation was the great genetic differences between geographically distant Yi populations (SCY and GZY). We further conducted model-based ADMIXTURE with the same data and observed that SCY shared more high-altitude Tibetan-related ancestral components (red). In contrast, GZY shared more low-altitude southern Chinese-related ancestral components (dark blue), demonstrating the existence of genetic substructure among Yi populations.

**Figure 3.**
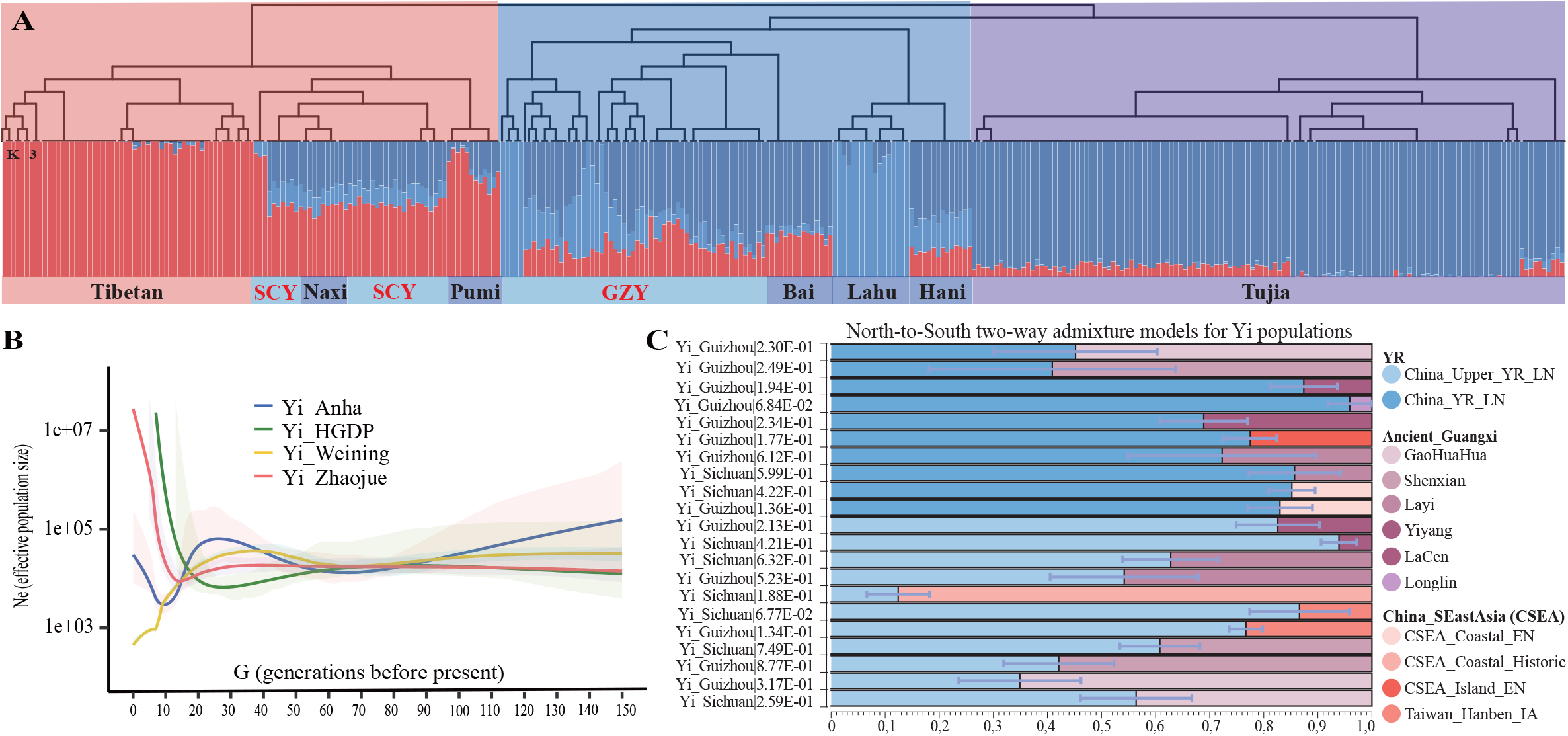
Fine-scale genetic analysis of SYC and GZY. **(A)** Explore the fine-scale genetic structure of TB populations based on shared haplotypes of 348 unrelated individuals. The Dendrogram in the top part shows the phylogenetic distribution by geographical pattern. ADMIXTURE in the bottom part corresponds to the individuals’ proportion of ancestral components of the TB population. From left to right are Ü-Tsang Tibetans, Yi, Naxi, Pumi, Bai, Lahu and Hani from TYC and YGP, and Tujia from the eastern lowlands. Notably, substructures exist among Yi. **(B)** The effective population size shown by Ne was used to determine whether a population bottleneck effect existed. GZY shows a decreasing trend from 30 generations ago to the present, suggesting that it has been experiencing population bottlenecks and supporting the possibility of genetic drift in GZY. In contrast, SCY shows the opposite trend, having experienced rapid population expansion in the last 10 generations. **(C)** Quantitative analysis of the proportion for SCY and GZY admixture by different north-south ancient sources was performed by qpAdm. We performed Two-way admixture models based on North-South ancestral sources identified by *f*_4_ tests. Northern Ancients: Yellow River Basin Ancients (YR, blue), including China_Upper_YR_LN and China_YR_LN. Southern Ancients: Ancient_Guangxi (purple), including GaoHuaHua, Shenxian, Layi, Yiyang, LaCen and Longlin. China_SEastAsia (red), including China_SEastAsia_Coastal_EN, China_SEastAsia_Coastal_Historic, China_SEastAsia_Island_EN and Taiwan_Hanben_IA.

We performed qpWave with worldwide representative populations as the outgroup sets to validate the genetic homozygosity and heterozygosity of geographically different Yi people. The values of p_rank0 focused on SCY and GZY confirmed their genetic differentiation (**Figure S11**). We sought to determine ancestral composition through qpAdm-based admixture models to explore the potential ancestral sources contributing to the identified genetic substructure. Two- and three-way admixture models showed the highest proportion of TB-related ancestral components. Notably, GZY had fewer northern Altaic-related components but more southern HM, AN and AA-related components than that in SCY (**Figure S12 and Tables S4-6**), indicating genetic heterogeneity within the same ethnic group in different areas due to genetic interactions and admixture with other diverse ethnic communities.

We then estimated effective population size (Ne) based on shared haplotypes to explore the demographic histories of geographically different Yi populations. The three Yi people from Sichuan shared a similar demographic history with rapid population expansion during the past ten generations. In contrast, GZY experienced a population bottleneck in the 30th generation, resulting in significantly low Ne (**Figure 3B**). We also performed runs of homozygosity (ROH) for TB populations and observed a trend of increasing ROH length from Ü-Tsang Tibetan and Yi people to Burmese-speaking populations (**Figure S13**), which illustrated less inbreeding within Ü-Tsang Tibetan groups and a moderate probability of inbreeding in Yi populations. Specifically, we found that SCY had higher ROH than GZY. We next estimated the admixture times of the inferred admixture events of focused Yi people (**Table S3**). Northern Han Chinese from Shaanxi and southern Chinese Zhuang populations met at 31.84-36.78±5.07-8.22 generations ago and formed the observed LD patterns of Yi people, which was congruent with the fact of historically documented strengthen-control by Qin and Han Dynasties ^51^.

### Genetic origin and complex admixture landscape of one underrepresented TB sub-lineage

The genetic history of northmost TB people of Tibetan and Sherpa and southmost Burmese speakers have been characterized in recent genomic work^13, 24, 25^. To fill the gap in the evolutionary history of middle TYC TB-speaking populations, we performed PCA based on the merged HO dataset with 45 ancient populations projected onto modern genetic coordinates (**Figure 1B**). we observed a west-to-east gradient including Tibetan and Sherpa populations in the TP, YR farmers, and TYC and YGP populations (Bai, Yi, Naxi, and Pumi) along PC1, which separated from Burmese-speaking groups in MSEA. This indicated that TB populations in China possessed closer genetic similarity with YR farmers than that of TB populations in MSEA. In addition, Ü-Tsang Tibetans living in the TP and Pumi from TYC formed a cluster with ancient individuals from Nepal (aMMD, ancient DNA data from Himalayan sites in the Mustang and Mannang districts) ^52^. ADMIXTURE results revealed different ancestral compositions among geographically differentiated TB populations. The red component appearing in aMMD groups occupied high proportions in Ü-Tsang Tibetans, while the pink component associated with YR farmers enriched in TYC and YGP populations (**Figure 2A**).

We carried out an *f*_4_-statistic in the form of *f*_4_(Northern-reference, Southern-reference; studied populations, Mbuti) based on the merged 1240K dataset to verify the observed pattern firmly (**Figure S14**). We selected geographically diverse ancient populations in the same time dimension as reference pairs, and Yi shared more alleles with ancient populations from North China (YR farmers) relative to ancient southern groups (**Figure S15**). Further, we performed asymmetric *f*_4_-statistic in the form of *f*_4_(Southern-reference, studied populations; YR, Mbuti) and observed significant negative scores, indicating that Yi harbored closer genetic connections with YR farmers than that of other ancient populations (**Figure S16**). Moreover, formal tests using *f*_4_(YR, studied populations; Southern-reference, Mbuti) provide direct evidence that Yi populations received additional gene flow from southern ancients (LaCen, GaoHuaHua and Taiwan_Hanben_IA) relative to their potential northern ancestry source, demonstrating that Yi populations were influenced by ancient southern groups (**Figure S17**). Moreover, to infer whether different ancient interactions happened between Yi populations and ancient groups, we conducted symmetry *f*_4_-analysis in the form of *f*_4_(SCY, GZY; ancient reference, Mbuti). We observed that GZY received more ancient interactions with southern Chinese than SCY, while SCY received more gene flows from North Chinese ancients than GZY (**Figure S18**).

We further conducted the two-way admixture models using qpAdm to investigate potential ancestral surrogates and corresponding proportions. SCY and GZY could be simulated as an admixture of ancient northern populations represented by YR ancients (0.124-0.939 and 0.349-0.959) and ancient southern populations represented by Guangxi ancients (0.061-0.939 and 0.041-0.959) or China_SEastAsia (0.134-0.876 and 0.17-0.233) (**Figure 3C and Table S7**). We also identified the North-South admixture pattern of Yi populations by identifying, describing and dating with GLOBETROTTER (**Table S8**). The available archaeological data suggest the hypothesis of the northern origin of Yi populations who migrated along TYC and intermixed with southern inhabitants ^53^.

### Shared and specific genetic adaptation signatures associated with population differentiation

To explore the adaptive signatures of meta-Yi populations, we first combined studied Yi populations as one targeted population and selected Mongolian_InnerM (IM) and CEU as the second and third outgroup reference populations. We conducted Population Branch Statistic (PBS) analysis, which identifies alleles exhibiting great changes in frequency in the studied population compared to reference outgroup populations. In the Yi-IM-CEU model (**Table S9**), we observed the strongest signal on ectodysplasin A receptor (*EDAR*, PBS = 0.36) located on chromosome 2 (rs260690), which encodes a transmembrane protein that is a receptor for the soluble ligand ectodysplasin A and modulates morphological change of teeth and increases the thickness of scalp hairs ^54^. When restricted to the top 0.1% of PBS values, two previously reported genes, *SLC24A5* and *ALDH9A1,* were identified, *SLC24A5* is associated with skin pigmentation ^55^ and *ALDH9A1* in the family of aldehyde dehydrogenases involved in alcohol metabolism ^56^. Missense variants for *CASC5, OAF, FAM214A, GAS8, HADHB* and *PUS10* were also found to be subjected to natural selection (**Table S9**). Strikingly, *CTNNA3*, located on chromosome 10, has an extremely high bio-signal (PBS = 0.32) and several mutation loci (rs2441727, rs2660024, rs1911341 and rs1911355), which encodes a protein (α-t-catenin) that plays a role in cell-cell adhesion of muscle cells ^57^. The local genomic region containing *CTNNA3* in zoomed view found *LRRTM3 gene* (leucine-rich repeat transmembrane neuronal 3) (**Figure 4A**), which is transcribed with the opposite direction to *CTNNA3*, acting as nested genes (a form of gene overlap) of *CTNNA3* to co-regulate tissue-specific expression ^57^. Functional annotation focused on *CTNNA3* via GeneORGANizer showed a correlation with cardiopulmonary function (**Figure 4B**). We further generated frequency spectrums for four loci of *CTNNA3* identified by PBS (**Figure 4C**) to detect the allelic divergence in the worldwide context. The derived allele of rs2441727(G), rs2660024(G), rs1911341(C), and rs1911355(C) have much higher frequencies in East Asians and island Southeast Asians than among other continent populations.

**Figure 4.**
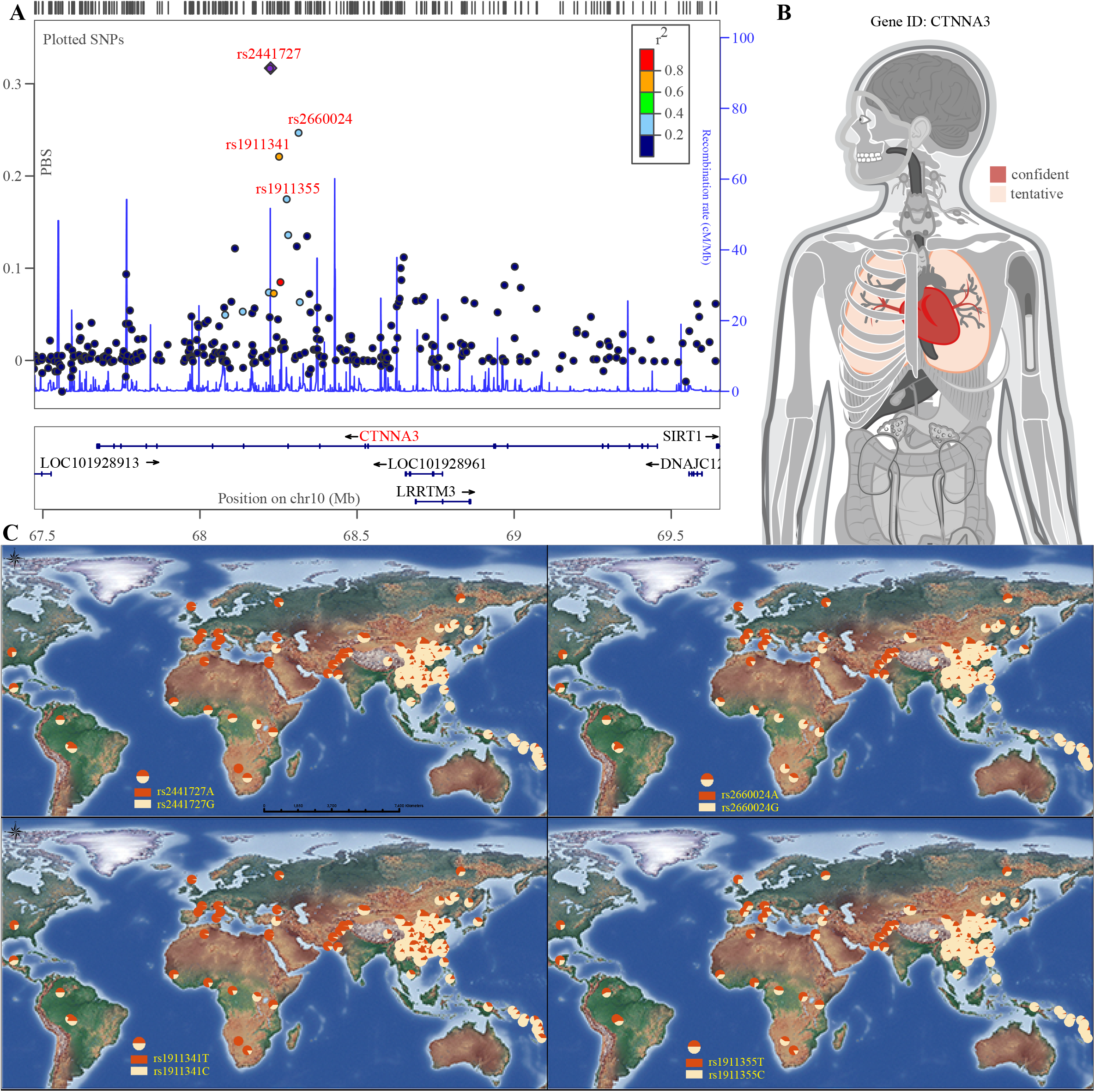
Nature selection of CTNNA3 found in Yi populations based on PBS. **(A)** The physical location of CTNNA3 genes with the higher and more frequent appearance in the TOP 0.01% based on PBS selection was observed by LocusZoom. **(B)** Functional annotation of CTNNA3 gene using GeneORGANizer. Cardiac function: confident (red) and tentative (pink) pulmonary function. **(C)** The Allele frequency spectrum was drawn for four loci (rs2441727, rs2660024, rs1911341, rs1911355) of the CTNNA3 gene based on 376 populations worldwide. Different distribution patterns of variation were shown for this gene in East and island Southeast Asia compared to European, America and Africa.

There was a striking stratification pattern across geographically distinct Yi populations. Here we calculated PBS for SCY and GZY, with IM and CEU regarded as ingroup and outgroup, respectively (**Table S10**). We found that putatively adaptive genes of *CSRP1, DGKB, NGF, PTPRT* and *HS6ST3* were shared between two Yi groups (**Table S11**), which jointly mediate in Cell Type Signature (GO: M39070). To investigate population-specific signatures of adaptation, we searched for signals specific to one of these two Yi populations (orange) (**Figure S19**). Most prominently, *SLC29A3, CACNA1C, SYNE2, PFKP* and *IRF5* were only observed in SCY (**Figure 5A**). The gene *SLC29A3*, which is involved in skin pigmentation ^58^, *CACNA1C*, *SYNE2* and *IRF5* are bio-functionally relevant genes and *PFKP* plays critical roles in regulating glycolysis ^59–62^. While *CGNL1, JAZF1-AS1, MACROD2, PEX11A* and *FAM189A1* were specific to GZY (**Figure 5B**), and *PEX11A* may be relevant in response to external stimuli ^63^. Moreover, Gene Ontology (GO) and Kyoto Encyclopedia of Genes and Genomes (KEGG) pathway analysis by Metascape demonstrated that the selected top 0.1% genes revealed the relevant pathways on adaptive physiological traits in Yi populations. We observed that several pathways enriched in SCY were associated with response to stimulus (GO:0050896), biological regulation (GO:0065007) and positive regulation of biological process (GO:0048518). In contrast, GZY harbored pathways regarding the metabolic process (GO:0008152), negative regulation of biological process (GO:0048519) and immune system process (GO:0002376) (**Figure S20**). We next constructed protein-protein interaction networks and found that SCY showed the most significant lgP values in ECM-receptor interaction (hsa04512), human papillomavirus infection (hsa05165) and retrograde neurotrophin signaling (R-HSA-177504). As for GZY, the most significant lgP values were in cellular component morphogenesis (GO:0032989), cell-cell adhesion (GO:0098609) and cell adhesion molecules (hsa04514) (**Table S12**). Collectively, we illuminated the shared and specific adaptive genetic signatures across geographically diverse Yi populations.

**Figure 5.**
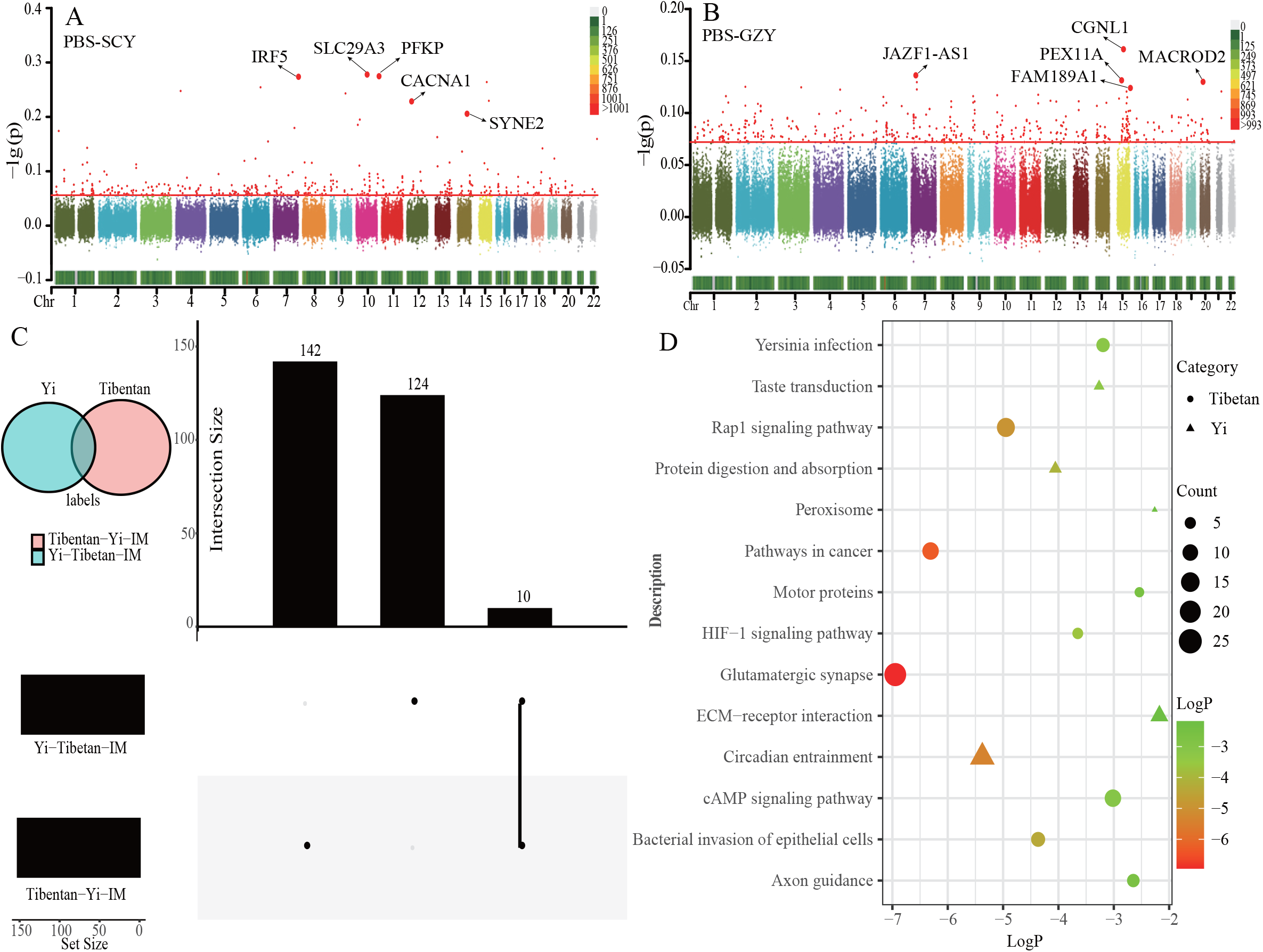
Local adaption identified within-ethnic and between-ethnic. **(A-B)** Differences in the highest biological adaptation signals for SCY and GZY under different geographical patterns, expressed in Manhattan plots, demonstrate local adaptation within-ethnic. **(C)** Wayne plots comparing specific PBS for Tibetan in extreme environments and Yi at middle altitude revealed 142 genes specific to Tibetan versus 124 genes specific to Yi. Evidence for local adaptation between-ethnic. **(D)** Bubble plots based on KEGG after enrichment of Tibetan and Yi. The category is sorted by ethnics, Tibetan (circles), and Yi (triangles). The number of genes in the enrichment pathway determines the shape’s size. LogP values represent the strength of the pathway signal, strong (red), and weak (green).

Tibetans living in high-altitude regions (TP) have developed a series of traits to adapt to harsh environments, and Yi populations in middle-altitude areas (TYC and YGP) may have evolved a suite of traits to cope with the regional environment. To explore the status of the shared and specific putative selected variants and genes, we calculated PBS for Yi and Tibetan populations, with Tibetan and Yi as second groups, respectively, and IM regarded as the third group (Yi-Tibetan-IM and Tibetan-Yi-IM) (**Figure 5C and Table S13**). Consistent with previous findings ^4^, several extremely high levels of high-altitude adaptation signals represented by *EPAS1* (PBS = 0.74), *EGLN* (PBS = 0.23), and *DISC1* (PBS = 0.19) were identified in Tibetan people. We conducted KEGG enrichment and revealed these genes’ polygenic and pleiotropic effects, which jointly affect the hypoxia-induced factor 1 signal pathway (HIF-1). Only *EPAS1* (PBS=0.09) was found in Yi populations with weak signals. We also performed KEGG enrichment for Yi populations and identified various pathways involved in circadian entrainment (hsa04713), ECM-receptor interaction (hsa04512), and protein digestion and absorption (hsa04974) (**Figure 5D**). We observed significant differences in population-specific or shared adaptive signatures between Tibetan populations living on the TP and Yi populations in the middle/low-altitude regions.

## DISCUSSIONS

Systematic sampling biases remain in human genomics studies of ethnically diverse minorities, which may introduce the missing diversity of human populations ^47^. Recent human pangenome projects focused on worldwide populations (Human Pangenome Reference Consortium, HPRC) and Chinese minorities (Chinese Pangenome Consortium, CPC) also highlighted that well-designed sampling strategies could provide new insights into human genetic diversity and patterns of categories of human variations and understanding of the biological consequence of newly-identified genetic variants ^64, 65^. This study presented an entire landscape of the genetic structure of geographically high-coverage TB people based on the genome-wide SNP data from the newly-generated and publicly collected datasets ^3, 23, 24^. Furthermore, we revealed that geographically dispersed TB groups were genetically different from each other. In contrast with previous genetic conclusions that TB people derived ancestry from ancient northern East Asian-related populations, our findings supported that geographically different TB people harbored different genetic affinities with northern and southern East Asians. The northern TB people shared more ancestry with ancient northern East Asians related to middle and late Neolithic millet farmers and modern Sinitic people, and the southern one obtained additional ancestral influence from HM/TK-related ancestral sources. We comprehensively illuminated the identified language-related population stratification via multiple statistical methodologies, including PCA, Fst, ADMIXTURE, outgroup-*f*_3_ and IBD (**Figures S1-7**). We found a geography-related genetic distinction, which divided TB people into Tibetan-related highlanders (Ü-Tsang Tibetans, Sherpa and Tamang), middle-altitude groups from TYC and YGP (Qiang, Yi, Bai, Naxi, Tibetan_Yajiang), as well as Burmese-speaking people from MSEA (**Figure 1**). Generally, our observation agreed with previous genetic studies, which mainly focused on frequent interactions among geographically close minorities ^5, 24, 27^. We also provided differentiated language-related population structure using ADMIXTURE, *f*_4_(GZY, SCY; Reference, Mbuti) and TreeMix. We proposed that this genetic differentiation could be ascribed to the regionally differentiated admixture with linguistically different ethnic groups (**Figures 2 and S8-9**). The minorities speaking the Yi language (SCY) and Tibetic language (Qiang, Tibetan subgroups and Kham Tibetan) living in the TYC area exhibited a genetically close relationship with highland Tibetans (Ü-Tsang Tibetans and Sherpa), revealing that they possessed similar patterns of cultural and geographical affinity ^52, 66^. We also observed the differentiation of ancestry profiles among Ü-Tsang/Ando/Kham/Subgroups Tibetans (**Figures S8 and S10**). A recent study found that Ando Tibetans in Northwest China harbored genetic influence from western Eurasians, and Kham Tibetans in Southwest China received strong gene flow from ancestral southern East Asians compared to Ü-Tsang Tibetans ^6^. Besides we also identified the differentiated genetic patterns in geographically different TYC people. The GZY, Bai, and Lahu living in the YGP had extensive genetic admixture with TK, AA and HM-related ancestral people ^27, 67^. The Burmese-speaking groups (Karen, Lisu, Cong, and Sila) from MSEA appeared to be related to AA and TK-speaking populations ^23, 24^.

We further reconstructed demographic histories to illuminate the influence of differentiated admixture processes on the observed patterns of genetic diversity and ancestral composition. Previous genetic studies based on WGS data of Uyghur have illuminated that admixture between previously isolated populations could contribute to higher genetic diversity, deleterious disease burden, relative homogeneous genetic structure and differentiated biological adaptation of admixed populations than their source populations ^68^. We not only observed genetic differences between geographically different TB people and reported the fine-scale genetic differences among geographically different Yi people in the TYC. We observed apparent genetic substructure and genetic heterogeneity between SCY and GZY (**Figures 3A and S11**), who experienced local admixture events under prolonged demographic history. Previous work showed that Yi had a medium-ROH value and a larger number of heterozygous site results among TB people ^5^, indicating they were heavily admixed and had low genetic drift. Our observation from Ne estimates based on shared IBD patterns suggested that GYZ experienced prolonged population bottlenecks 20-30 generations ago. In contrast, SYC experienced population expansion during this period (**Figure 3B**), possibly owing to the human migration from Guizhou to Sichuan caused by the Qing Dynasty’s policy of bureaucratization of native officers ^51^. In addition, we found that SCY possessed genetic homogeneity with Bai from Yunnan province based on the qpWave results (**Figure S11**). The historical record showed that their ancestry established Nanzhao Kingdom in the 8th century AD and had the same social status compared to other TB minorities who were under control ^69^.

Paleogenetic data provide a powerful source for tracing human origin. Combined with the evidence from linguistics and archaeology, the northern YR origin hypothesis of TB populations has been supported ^11, 66^. Our results reconfirmed this hypothesis and further identified the admixture pattern between YR millet farmer-related ancestry and southern aborigines by PAC, ADMIXTURE, *f*_4_-statistics, qpAdm and GLOBETROTTER (**Figures 1-3, S14-17 and Tables S6-7**). Coinciding with the traditional recordings and similar cultural elements, their ancestors moved southwards through TYC due to famine and war during the historical period and admixed with southern indigenous inhabitants ^53^. In particular, we also observed genetic differentiation between geographically different Yi populations in which GZY received more gene flows from Southern ancients (LaCen, GaoHuaHua and Hanben_IA) than SCY (**Figure S18**). Taking advantage of the genome-wide SNP data newly reported for Yi people and publicly available HGDP genomic resources, we presented the comprehensively evolutive trajectory of Yi populations. As a supplementary of previous studies that trace TP Tibetans’ multi-wave colonization and permanent settlement [9, 18], our results conjointly provided the genetic evidence of the southward footprint of TB-speaking populations. One is represented by Tibetic speakers who traversed the TP from its northeastern fringe, and the other is represented by Yi-speaking populations southward along the TYC ^70, 71^. The genetic, historical and linguistic proofs demonstrated that although TB speakers had the same genetic origin, they showed distinct patterns of genetic admixture since the split from Han Chinese ^46^. Recent ancient DNA from the late Neolithic Gaoshancheng site in Chengdu Plateau and Bronze Age Haimenkou site in YGP also identified a strong genetic connection between ancient YR millet farmers and TYC-related ancient people ^72^. The ancient genetic connections between northern and southern TYC people and similar cultural elements were also evidenced in our work, suggesting both demic and cultural diffusion hypotheses have contributed to the complex gene pool of Yi people. However, as the lack of ancient DNA data from the Sanxingdui and Jinsha sites, we have no clues to illuminate the direct genetic relationship between the Yi people and the ancient Sanxingdui and Jinsha people.

It has been suggested that genomic diversity is determined by genetic and environmental factors ^73^. Concerning extremely extraordinary adaptive signals shared in Tibetan people, previous studies concentrated on the Darwinian selection of hypoxic adaption introgressed (*EPAS1*) from archaic Denisovan people ^4, 74^ and genetic adaptation of skin pigmentation (*GNPAT*) in highland groups ^75^. We searched for local adaption signals in Yi populations by scanning putative adaptive genes via PBS (Yi-IM-CEU trio) (**Table S8**). *EDAR* and *SLC24A5*, responsible for skin pigmentation-related genes, were previously reported as candidate genes favored by natural selection in human populations ^54, 55^. Simultaneously, the *ALDH9A1* gene, involved in liver alcohol metabolism, was possibly caused by long-term lifestyle habits [54]. Intriguingly, we also identified the *CTNNA3* genes (frequent appearance and high PBS value) (**Figure 4A**), which encodes alpha-T-catenin and regulates the biological function of cell-cell adhesion ^57^. *CTNNA3* is the fifth largest gene in the human genome and is located on the common fragile site *FRA10D*, an unstable and important locus for screening many diseases. Therefore, it has an important biological meaning ^57^. Relevant study on Tibetan adaptive genetic phenotypes has involved numerous cardiovascular diseases, such as arrhythmogenic right ventricular cardiomyopathy (ARVC) and dilated cardiomyopathy (DCM) ^76^. The *CTNNA3* found in Yi populations was also confirmed by animal experiments that expression of this gene leads to cardiomyopathy (**Figure 4B**) ^57^. The association between genes and phenotypes demonstrates the significance of local adaptation research for ethnic groups. The construction of ethnic-specific databases will help us to achieve the ultimate goal of personalized health management and accurate disease risk prediction.

Further investigation based on PBS results and functional enrichment assays found that SCY and GZY possessed differential adaptive signals (SCY-GZY-IM trio and GZY-SCY-IM trio) (**Figures 5A-B, S19-20 and Tables S9-11**). A similar result was observed in seeking the local distinction-adaption between Yi and Tibetan who live on the TP (Yi-Tibetan-IM trio and Tibetan-Yi-IM trio) (**Figures 5C-D and Tables S20**). Our tentative findings indicated genetic diversity among TB speakers, which may reflect the influence of geographical barriers and existing differentiated evolutionary processes. To comprehensively confirm our findings, high-coverage sequencing data and representative ethnic genomic data need to be generated in future studies. Recently, multiple studies were performed to explore the human variants associated with their habits, disease risk, environments, or even ethnic cultures. Yang et al. speculated that the descendants of the Di-Qiang lineages were likely under habit-related adaptive evolution of hypertension ^27^. Analyzing the fine-scale genetic diversity of TB people can be biomedically multidimensional informative, including diagnosing disease, selecting therapies and predicting treatment outcomes. Therefore, it is urgently required to construct a full-areas, high-density and representative database for TB populations as their ethnolinguistics and culture are rapidly losing.

## CONCLUSIONS

The genetic history and biological adaptation of high-altitude Tibeto-Burman (TB) people were complicated because the Tibetan Plateau (TP) and surrounding areas possess hypoxic, cold and high-UV subsistence conditions. However, the genetic investigation of differentiation processes of ethnolinguistically and geographically diverse TB people has not been exhaustive. To characterize a comprehensive landscape of genetic diversity and admixture history of geographically different TB people and dissect the differentiated adaptative features of highland and lowland TB people, we generated a meta-database consisting of the newly-generated and publicly available genome-wide SNP data of 500 individuals from 39 TB populations from East Asia and Southeast Asia. Population admixture models based on sharing alleles and haplotypes showed geography-related population genetic substructure. Geographically diverse TB people could be divided into the western highland group including Tibetan and Sherpa in the Ü-Tsang region, the central middle-altitude group consisting of Qiang, Bai, Yi, Naxi and Lahu populations from the Tibetan-Yi corridor and the Yunnan-Guizhou Plateau, and lowland Burmese-speaking ethnic groups from the Alluvial Plain and mainland Southeast Asia. TB people from the TP harbored dominant ancient Yellow River (YR) millet farmer ancestry, and lowland TB people received more genetic influence from ancient southern East Asians. We identified extensive ethnicity-related fine-scale substructures, in which genetic differentiation between geographically distant Yi and massive differentiated admixture events among geographically close but ethnically different populations were first deeply reported here. We illustrated a robust genetic connection between Yi populations and ancient YR people, supporting the Northern origin hypothesis of TB people. We identified differentiated biological adaptative signatures between highland Tibetans and lowland TB people and between geographically different Yi populations and found adaptative signals associated with the pairwise highly-differentiation variants. Geographically different TB people shared differentiated putative adaptive genes and pathways. Hypoxic adaptative signatures related to *EPAS1* and *EGLN* were confirmed only in Tibetans from TP, adaptation associated with the physical features and skin pigmentation (*EDAR* and *SLC24A5*), hepatic alcohol metabolism (*ALDH9A1*) and regulation of cell-cell adhesion of muscle cells (*CTNNA3)* existed in western Yi from Tibetan-Yi corridor, and immune or fat metabolism-related adaptative signatures were identified in eastern Yi from Yunnan-Guizhou Plateau. We provided comprehensive integrative genomic resources that offered insights into the fine-scale population substructure and biological adaptation of TB people. We identified differentiated genetic structures and adaptation signatures from the geography-related different admixture models.

## METHODS AND MATERIALS

### Sample collection, genotyping and quality control

We collected 47 saliva samples of Yi individuals from Anha (24) and Zhaojue (23) in Liangshan Yi Autonomous Prefecture of Sichuan Province, China (**Figure S1**). All volunteers were obtained with written informed consent. They claimed that they were indigenous residents of Yi for three or more generations and were unrelated within three generations. Our study followed the recommendations of the Declaration of Helsinki revised in 2000 ^77^ and the regulations of China’s Human Genetic Resources Administration (HGRAC). The relevant protocols in our project were reviewed and approved by the Medical Ethics Committee of West China Hospital, Sichuan University (2023-306). Forty-seven individuals first reported here were genotyped by the Illumina array. The genetic relatedness between sampled individuals was examined using PLINK v.1.90 ^78^ and King ^79^. We have estimated PI_HAT values with the PLINK ‘-genomic’ parameter. Coefficient estimates were further validated with the “--correlation -- ibs” parameter of King for individual pairs whose values were greater than 0.15. The final quality-control dataset contained ∼516,448 SNPs.

### Dataset merging

To provide a representative genetic investigation of TB ethnic groups, we included previously published and publicly available data from 500 unrelated individuals from 39 populations ^3, 23, 24^, including Ü-Tsang Tibetans (Tibetan_Shigatse, Tibetan_Lhasa and Tibetan_Chamdo) from the core-Tibet region, Ando Tibetans (Tibetan_Gangcha, Tibetan_Gannan and Tibetan_Xunhua) resided in Gansu-Qinghai region, Kham Tibetans (Tibentan_Xinglong, Tibentan_Yajiang and Tibentan_Yunnan), Tibetan subgroups (Pumi and Naxi), Yi (SCY and GZY), Qiang, Bai, Lahu and Hani living in Southwest China, Tujia in Eastern lowland, Burmese-speaking ethnic groups (Cong, HaNhi, Karen, Lisu and Lolo) in MSEA (**Figure S1 and Table S1**). We mainly generated three merged datasets: (1) To retain more SNPs for fine-scale genetic structure and natural selection analysis, we used the Illumina_HGDP high-density dataset (including 460,678 SNPs) that merged with WGS data ^80, 81^. (2) To explore the gene flow events of ancestral interactions and the interactions among modern populations, we used the Illumina_1240K dataset comprising 146,802 SNPs ^82^. (3) To clearly represent the genetic diversity of modern people and ancient reference populations, we used the Illumina_HO dataset, including 56,814 SNPs, which was characterized by a large range of population sizes. We retrieved publicly available datasets from HGDP ^83^ and Oceania genomic resources ^49^, AADR (HO and 1240K datasets) from David Reich Lab (https://reich.hms.harvard.edu/datasets).

### Principal Component Analysis

Principal component analysis was performed using the smartpca program of the EIGENSOFT package ^84^, which aims to investigate the clustering pattern between studied and reference populations. We conducted a clustering analysis based on the Illumina_HGDP dataset, including 203 populations at the world-scale. The clustering pattern at the East Asian-scale and the TB-scale were shown based on the Illumina_HO dataset. The parameters were set by numoutlieriter: 0 and lsqproject: YES. Modern populations were first positioned on the plane with coordinates formed by PC1 and PC2, and then 45 ancient populations were projected on the aforementioned two-dimensional plot.

### ADMIXTURE Analysis

We first pruned out SNPs with linkage disequilibrium (LD) via PLINK v.1.9 41, with parameters set of -indep-pairwise 200 25 0.4. Model-based ADMIXTURE ^85^ analysis was carried out based on a maximum likelihood clustering algorithm, and we ran admixture models based on the Illumina_HO dataset with predefined ancestral sources (K values) ranging from 2 to 20, using different random seeds in the bootstrap sequence. The best-fitted model was selected by calculating the cross-validation (CV) error.

### Genetic difference analyses

We calculated the pairwise fixation indexes (Fst) using PLINK v.1.90 ^13^ to estimate the genetic differences between studied and reference populations at different scales. In addition, NJ trees were constructed via MEGA based on the pairwise Fst values to investigate the phylogenetic relationship.

### F-Statistics

We used the qp3Pop software in ADMIXTOOLS ^82, 86^ to calculate the outgroup-*f*_3_-statistics in the form of *f*_3_(X, Y; Outgroup), which was used to estimate genetic drift sharing among X and Y populations further to demonstrate genetic affinity between studied populations and reference populations if the larger *f*_3_ value was observed among X and Y, the more shared genetic drift. Here, the X represents Yi populations, the Y represents the other TB groups, and Mbuti from central Africa is used as the outgroup. We generated an NJ tree based on the 1-outgroup-*f*_3_ values to observe the genetic affinity within TB populations.

We used the qpDstat software in ADMIXTOOLS to perform comparisons of four groups with parameters of *f*_4_: YES. We compared shared alleles based on the Illumina_HO dataset. We then filtered out representative or characteristic populations and compared them using the Illumina_1240K dataset. There are three general forms of model construction, the first of which is the symmetrical *f*_4_, expressed as (Studied1, Studied2; Reference populations, outgroup), to examine whether two studied populations form a clade compared to reference populations. The asymmetrical *f*_4_-statistics in the form of *f*_4_(Reference1, Studied populations; Reference2, outgroup) and the affinity *f*_4_-statistics in the form of *f*_4_(Reference1, Reference2; Studied populations, outgroup) were used to test whether studied populations shared more alleles with Reference1 or Reference2. Mbuti is commonly used as an outgroup population.

### Effective population size and runs of homozygosity

We used SHAPEIT v2.r90 ^87^ for haplotype phasing to evaluate the shared IBD segments within and between Yi populations. IBDNe v23Apr20 ^88^ was used to extrapolate the variation of effective population size (Ne) between 1 and 150 generations. Refined IBD ^89^ lengths were divided into three-stage datasets from 1 to 5, 5 to 10 and above 10cM. We tested ROH using sliding windows of 5Mb for studied populations and TB populations using PLINK 1.9 ^78^ to measure the inbreeding relationships. To further elaborate the information contained in ROH, we categorized ROH lengths into <1, 1-5 and >5Mb levels. The average length of ROH for each individual was calculated as the total length of ROH/the number of ROH.

### Maximum Likelihood Tree

We generated a maximum likelihood tree representing gene flow events from 0 to 7 based on allele frequencies using TreeMix v.1.13 ^90^, with parameters of -se, -bootstrap (100 bootstrap replicate), -k ( 500 group SNPs to account for the LD) and -global, to explore the phylogenetic relationship between studied populations and surrounding ethnic populations.

### Admixture modeling

A pairwise qpWave test was performed to determine whether Yi populations were genetically homogeneous, which was judged by p-values calculated by ADMIXTOOLS ^82^, p-values > 0.05 suggested that the population pairs were genetically homogeneous. In contrast, p-values < 0.05 suggested that populations were genetically heterogeneous. We calculated the correspondingly ancestral proportions of Yi populations using qpAdm ^82^ packages in the ADMXITTOOLS. SCY and GZY were used as tested populations, respectively. We used the following populations as the core outgroups: Mbuti, Iran_GanjDareh_N, Italy_North_Villabruna_HG, Ami, Mixe, Onge, Papuan and Russia_Ust_Ishim_HG. When Cambodians were used as the additional outgroup of present-day people, source populations including Altaic people (Daur, Oroqen, Mongolian, Xibo and Hezhen) were the northern ancestral sources. TB (Hani_Honghe, Lahu_Lancang and Bai_Heqing), HM (Yao_Yaoai and Yao_Gulei), AA (Wa_Cangyuan and Blang_Menghai), AN-speaking populations (Paiwan_Taiwan and Atayal_Taiwan) were adopted as the southern ancestral source, respectively. When Australian.DG as the additional outgroup of ancient people, source populations including YR people (China_YR_LN and China_Upper_YR_LN) were used as the northern ancestral source, Guangxi ancients (GaoHuahua, Shenxian, Layi, Longlin and Yiyang) and southeastern coastal ancients (Hanben_IA, China_SEastAsia_Island_EN, China_SEastAsia_Coastal_EN and China_SEastAsia_Coastal_Historic) were used as the southern ancestral source, respectively. The best-fitting model of qpAdm obeyed three principles as follows: (1) The value of ancestry proportion > 0, (2) the P value > 0.05, (3) the standard error larger than the minimum mixture proportion.

### Estimating Admixture time with ALDER

We evaluated demographic admixture events based on LD. We used ALDER v1.03 to observe the diminishing of LD and estimate the possible admixture time in Yi populations ^91^. Numerous modern people from East Asia were included in the simulations as potential ancestral populations, with all possible source combinations tested. Parameters were set of mindis: 0.005, jackknife: YES.

### FineSTRUCTURE

To dissect whether substructure exists among Yi populations from different areas, we performed a painting of all individuals based on the Illumina_HGDP dataset using ChromoPainter v2 ^92^, which generated a Dendrogram tree. We also presented the fine-scale genetic structure based on haplotype information using fineSTRUCTURE 4.1.0 ^92^.

### Natural selection signal index

PBS was used to identify natural selection signals in our study. PBS was calculated with the formula = (T_AB_ + T_AC_ - T_BC_)/2, T = -lg(1-Fst), and A was the targeted population, B and C were the ingroup and outgroup, respectively. To explore the adaptive signals of natural selection within Yi populations, we combined Yi populations and used IM as the ingroup and CEU as the outgroup to investigate the selection characteristics of studied populations. Based on the inferred demographic models, we conducted a Tibetan_Tibet-IM-CEU trio to compare the genetic patterns between Tibetan located in the TP and Yi. We next performed SCY/GZY-specific PBS with respect to genetic differences between SCY and GZY.

## DATA AVAILABILITY

The allele frequency data derived from human samples have been deposited in the National Omics Data Encyclopedia (NODE, http://www.biosino.org/node) and can be accessed with accession number (OEPXXXXXX, available after publication). The access and use of the data shall comply with the regulations of the People’s Republic of China on the administration of human genetic resources. Requests for access to data can be directed to Guanglin He.

## ACKNOWLEDGMENTS

This work was supported by grants from the National Natural Science Foundation of China (82202078). We thank Prof. Etienne Patin and Prof. Lluis Quintana-Murci from the Human Evolutionary Genetics Unit of the Institute Pasteur for sharing the high-coverage genomes of 317 individuals from the Pacific region. We thank Prof. Mark Stoneking, Prof. Dang Liu at Max Planck Institute for Evolutionary Anthropology, and Prof. Wibhu Kutanan at Khon Kaen University for sharing genome-wide SNP data from Vietnam, Thailand, and Laos.

## AUTHOR CONTRIBUTIONS

G.H., L.Y. and M.W. conceived and supervised the project. Y.S., G.H., M.W. and J.Z. collected the samples. G.H., M.W. and J.C. performed the extraction of the genomic DNA and coordinated the genome sequencing. Y.S., Q.S., S.D., X.D., Z.W., Y.Z., X.L., X.J., X.H., Y.L., H.S., Y.H., Y.C., J.Z., S.N., J.Y., R.T., C.W., C.L., X.D., M.W., G.H., L.H. and L.Y. performed population genetic analysis. Y.S., M.W. and G.H. drafted the manuscript. L.Y., X.D., M.W. and G.H. revised the manuscript.

### Ethics approval and consent to participate

The Medical Ethics Committees of West China Hospital of Sichuan University (2023-306) approved this study. This study was conducted in accordance with the principles of the Helsinki Declaration.

## Reference

1. van Driem, G. (2002). Tibeto-Burman replaces Indo-Chinese in the 1990s: review of a decade of scholarship. Lingua 112, 79–102. https://doi.org/10.1016/S0024-3841(01)00039-0.

2. Chen, F., Welker, F., Shen, C.C., Bailey, S.E., Bergmann, I., Davis, S., Xia, H., Wang, H., Fischer, R., Freidline, S.E., et al. (2019). A late Middle Pleistocene Denisovan mandible from the Tibetan Plateau. Nature 569, 409–412. 10.1038/s41586-019-1139-x.

3. Wang, C.C., Yeh, H.Y., Popov, A.N., Zhang, H.Q., Matsumura, H., Sirak, K., Cheronet, O., Kovalev, A., Rohland, N., Kim, A.M., et al. (2021). Genomic insights into the formation of human populations in East Asia. Nature 591, 413–419. 10.1038/s41586-021-03336-2.

4. Zheng, W., He, Y., Guo, Y., Yue, T., Zhang, H., Li, J., Zhou, B., Zeng, X., Li, L., Wang, B., et al. (2023). Large-scale genome sequencing redefines the genetic footprints of high-altitude adaptation in Tibetans. Genome Biol 24, 73. 10.1186/s13059-023-02912-1.

5. Zhang, Z., Zhang, Y., Wang, Y., Zhao, Z., Yang, M., Zhang, L., Zhou, B., Xu, B., Zhang, H., Chen, T., et al. (2022). The Tibetan-Yi region is both a corridor and a barrier for human gene flow. Cell Rep 39, 110720. 10.1016/j.celrep.2022.110720.

6. He, G., Wang, M., Zou, X., Chen, P., Wang, Z., Liu, Y., Yao, H., Wei, L.H., Tang, R., Wang, C.C., and Yeh, H.Y. (2021). Peopling History of the Tibetan Plateau and Multiple Waves of Admixture of Tibetans Inferred From Both Ancient and Modern Genome-Wide Data. Front Genet 12, 725243. 10.3389/fgene.2021.725243.

7. Yu, X., and Li, H. (2021). Origin of ethnic groups, linguistic families, and civilizations in China viewed from the Y chromosome. Molecular genetics and genomics: MGG 296, 783–797. 10.1007/s00438-021-01794-x.

8. Zhang, D., Shen, X., Cheng, T., Xia, H., Liu, W., Gao, X., and Chen, F. (2020). New advances in the study of prehistoric human activity on the Tibetan Plateau. Chinese Science Bulletin 65, 475–482. 10.1360/tb-2019-0382.

9. Liu, L., Chen, J., Wang, J., Zhao, Y., and Chen, X. (2022). Archaeological evidence for initial migration of Neolithic Proto Sino-Tibetan speakers from Yellow River valley to Tibetan Plateau. Proc Natl Acad Sci U S A 119, e2212006119. 10.1073/pnas.2212006119.

10. Sagart, L., Jacques, G., Lai, Y., Ryder, R.J., Thouzeau, V., Greenhill, S.J., and List, J.M. (2019). Dated language phylogenies shed light on the ancestry of Sino-Tibetan. Proc Natl Acad Sci U S A 116, 10317–10322. 10.1073/pnas.1817972116.

11. Zhang, M., Yan, S., Pan, W., and Jin, L. (2019). Phylogenetic evidence for Sino-Tibetan origin in northern China in the Late Neolithic. Nature 569, 112–115. 10.1038/s41586-019-1153-z.

12. Zhang, C., Lu, Y., Feng, Q., Wang, X., Lou, H., Liu, J., Ning, Z., Yuan, K., Wang, Y., Zhou, Y., et al. (2017). Differentiated demographic histories and local adaptations between Sherpas and Tibetans. Genome Biol 18, 115. 10.1186/s13059-017-1242-y.

13. Lu, D., Lou, H., Yuan, K., Wang, X., Wang, Y., Zhang, C., Lu, Y., Yang, X., Deng, L., Zhou, Y., et al. (2016). Ancestral Origins and Genetic History of Tibetan Highlanders. Am J Hum Genet 99, 580–594. 10.1016/j.ajhg.2016.07.002.

14. Liu, Y., Wang, M., Chen, P., Wang, Z., Liu, J., Yao, L., Wang, F., Tang, R., Zou, X., and He, G. (2021). Combined Low-/High-Density Modern and Ancient Genome-Wide Data Document Genomic Admixture History of High-Altitude East Asians. Front Genet 12, 582357. 10.3389/fgene.2021.582357.

15. Wang, M., Du, W., Tang, R., Liu, Y., Zou, X., Yuan, D., Wang, Z., Liu, J., Guo, J., Yang, X., et al. (2022). Genomic history and forensic characteristics of Sherpa highlanders on the Tibetan Plateau inferred from high-resolution InDel panel and genome-wide SNPs. Forensic Sci Int Genet 56, 102633. 10.1016/j.fsigen.2021.102633.

16. Zou, X., Wang, Z., He, G., Wang, M., Su, Y., Liu, J., Chen, P., Wang, S., Gao, B., Li, Z., and Hou, Y. (2018). Population Genetic Diversity and Phylogenetic Characteristics for High-Altitude Adaptive Kham Tibetan Revealed by DNATyper(TM) 19 Amplification System. Front Genet 9, 630. 10.3389/fgene.2018.00630.

17. He, G., Wang, Z., Wang, M., Luo, T., Liu, J., Zhou, Y., Gao, B., and Hou, Y. (2018). Forensic ancestry analysis in two Chinese minority populations using massively parallel sequencing of 165 ancestry-informative SNPs. Electrophoresis 39, 2732–2742. 10.1002/elps.201800019.

18. Su, B., Xiao, C., Deka, R., Seielstad, M.T., Kangwanpong, D., Xiao, J., Lu, D., Underhill, P., Cavalli-Sforza, L., Chakraborty, R., and Jin, L. (2000). Y chromosome haplotypes reveal prehistorical migrations to the Himalayas. Hum Genet 107, 582–590. 10.1007/s004390000406.

19. Basnet, R., Rai, N., Tamang, R., Awasthi, N.P., Pradhan, I., Parajuli, P., Kashyap, D., Reddy, A.G., Chaubey, G., Das Manandhar, K., et al. (2023). The matrilineal ancestry of Nepali populations. Hum Genet 142, 167–180. 10.1007/s00439-022-02488-z.

20. Wang, L.X., Lu, Y., Zhang, C., Wei, L.H., Yan, S., Huang, Y.Z., Wang, C.C., Mallick, S., Wen, S.Q., Jin, L., et al. (2018). Reconstruction of Y-chromosome phylogeny reveals two neolithic expansions of Tibeto-Burman populations. Mol Genet Genomics 293, 1293–1300. 10.1007/s00438-018-1461-2.

21. Wen, B., Xie, X., Gao, S., Li, H., Shi, H., Song, X., Qian, T., Xiao, C., Jin, J., Su, B., et al. (2004). Analyses of genetic structure of Tibeto-Burman populations reveals sex-biased admixture in southern Tibeto-Burmans. Am J Hum Genet 74, 856–865. 10.1086/386292.

22. Zhao, M., Kong, Q.P., Wang, H.W., Peng, M.S., Xie, X.D., Wang, W.Z., Jiayang, Duan, J.G., Cai, M.C., Zhao, S.N., et al. (2009). Mitochondrial genome evidence reveals successful Late Paleolithic settlement on the Tibetan Plateau. Proc Natl Acad Sci U S A 106, 21230–21235. 10.1073/pnas.0907844106.

23. Kutanan, W., Kampuansai, J., Srikummool, M., Brunelli, A., Ghirotto, S., Arias, L., Macholdt, E., Hubner, A., Schroder, R., and Stoneking, M. (2019). Contrasting Paternal and Maternal Genetic Histories of Thai and Lao Populations. Mol Biol Evol 36, 1490–1506. 10.1093/molbev/msz083.

24. Kutanan, W., Liu, D., Kampuansai, J., Srikummool, M., Srithawong, S., Shoocongdej, R., Sangkhano, S., Ruangchai, S., Pittayaporn, P., Arias, L., and Stoneking, M. (2021). Reconstructing the Human Genetic History of Mainland Southeast Asia: Insights from Genome-Wide Data from Thailand and Laos. Mol Biol Evol 38, 3459–3477. 10.1093/molbev/msab124.

25. Liu, D., Duong, N.T., Ton, N.D., Van Phong, N., Pakendorf, B., Van Hai, N., and Stoneking, M. (2020). Extensive Ethnolinguistic Diversity in Vietnam Reflects Multiple Sources of Genetic Diversity. Mol Biol Evol 37, 2503–2519. 10.1093/molbev/msaa099.

26. Lipson, M., Cheronet, O., Mallick, S., Rohland, N., Oxenham, M., Pietrusewsky, M., Pryce, T.O., Willis, A., Matsumura, H., Buckley, H., et al. (2018). Ancient genomes document multiple waves of migration in Southeast Asian prehistory. Science 361, 92–95. 10.1126/science.aat3188.

27. Yang, Z., Chen, H., Lu, Y., Gao, Y., Sun, H., Wang, J., Jin, L., Chu, J., and Xu, S. (2022). Genetic evidence of tri-genealogy hypothesis on the origin of ethnic minorities in Yunnan. BMC Biol 20, 166. 10.1186/s12915-022-01367-3.

28. Yao, H.B., Tang, S., Yao, X., Yeh, H.Y., Zhang, W., Xie, Z., Du, Q., Ma, L., Wei, S., Gong, X., et al. (2017). The genetic admixture in Tibetan-Yi Corridor. Am J Phys Anthropol 164, 522–532. 10.1002/ajpa.23291.

29. Zou, X., He, G., Wang, M., Huo, L., Chen, X., Liu, J., Wang, S., Ye, Z., Wang, F., Wang, Z., and Hou, Y. (2020). Genetic diversity and phylogenetic structure of four Tibeto-Burman-speaking populations in Tibetan-Yi corridor revealed by insertion/deletion polymorphisms. Mol Genet Genomic Med 8, e1140. 10.1002/mgg3.1140.

30. He, G., Chen, P., Zou, X., Chen, X., Song, F., Yan, J., and Hou, Y. (2017). Genetic polymorphism investigation of the Chinese Yi minority using PowerPlex(R) Y23 STR amplification system. Int J Legal Med 131, 663–666. 10.1007/s00414-017-1537-2.

31. Beall, C.M., Cavalleri, G.L., Deng, L., Elston, R.C., Gao, Y., Knight, J., Li, C., Li, J.C., Liang, Y., McCormack, M., et al. (2010). Natural selection on EPAS1 (HIF2alpha) associated with low hemoglobin concentration in Tibetan highlanders. Proc Natl Acad Sci U S A 107, 11459–11464. 10.1073/pnas.1002443107.

32. Jeong, C., Ozga, A.T., Witonsky, D.B., Malmstrom, H., Edlund, H., Hofman, C.A., Hagan, R.W., Jakobsson, M., Lewis, C.M., Aldenderfer, M.S., et al. (2016). Long-term genetic stability and a high-altitude East Asian origin for the peoples of the high valleys of the Himalayan arc. Proc Natl Acad Sci U S A 113, 7485–7490. 10.1073/pnas.1520844113.

33. Simonson, T.S., Yang, Y., Huff, C.D., Yun, H., Qin, G., Witherspoon, D.J., Bai, Z., Lorenzo, F.R., Xing, J., Jorde, L.B., et al. (2010). Genetic evidence for high-altitude adaptation in Tibet. Science 329, 72–75. 10.1126/science.1189406.

34. Sirugo, G., Williams, S.M., and Tishkoff, S.A. (2019). The Missing Diversity in Human Genetic Studies. Cell 177, 26–31. 10.1016/j.cell.2019.02.048.

35. Zhang, P., Luo, H., Li, Y., Wang, Y., Wang, J., Zheng, Y., Niu, Y., Shi, Y., Zhou, H., Song, T., et al. (2021). NyuWa Genome resource: A deep whole-genome sequencing-based variation profile and reference panel for the Chinese population. Cell Rep 37, 110017. 10.1016/j.celrep.2021.110017.

36. Guanglin, H., Hongbing, Y., Qiuxia, S., Shuhan, D., Renkuan, T., Jing, C., Zhiyong, W., Yuntao, S., Xiangping, L., Shaomei, W., et al. (2023). Whole-genome sequencing of ethnolinguistic diverse northwestern Chinese Hexi Corridor people from the 10K_CPGDP project suggested the differentiated East-West genetic admixture along the Silk Road and their biological adaptations. bioRxiv, 2023.2002.2026.530053. 10.1101/2023.02.26.530053.

37. Cong, P.K., Bai, W.Y., Li, J.C., Yang, M.Y., Khederzadeh, S., Gai, S.R., Li, N., Liu, Y.H., Yu, S.H., Zhao, W.W., et al. (2022). Genomic analyses of 10,376 individuals in the Westlake BioBank for Chinese (WBBC) pilot project. Nat Commun 13, 2939. 10.1038/s41467-022-30526-x.

38. Cao, Y., Li, L., Xu, M., Feng, Z., Sun, X., Lu, J., Xu, Y., Du, P., Wang, T., Hu, R., et al. (2020). The ChinaMAP analytics of deep whole genome sequences in 10,588 individuals. Cell Res 30, 717–731. 10.1038/s41422-020-0322-9.

39. Jeong, C., Wang, K., Wilkin, S., Taylor, W.T.T., Miller, B.K., Bemmann, J.H., Stahl, R., Chiovelli, C., Knolle, F., Ulziibayar, S., et al. (2020). A Dynamic 6,000-Year Genetic History of Eurasia’s Eastern Steppe. Cell 183, 890–904 e829. 10.1016/j.cell.2020.10.015.

40. Mao, X., Zhang, H., Qiao, S., Liu, Y., Chang, F., Xie, P., Zhang, M., Wang, T., Li, M., Cao, P., et al. (2021). The deep population history of northern East Asia from the Late Pleistocene to the Holocene. Cell 184, 3256–3266 e3213. 10.1016/j.cell.2021.04.040.

41. Wang, T., Wang, W., Xie, G., Li, Z., Fan, X., Yang, Q., Wu, X., Cao, P., Liu, Y., Yang, R., et al. (2021). Human population history at the crossroads of East and Southeast Asia since 11,000 years ago. Cell 184, 3829–3841 e3821. 10.1016/j.cell.2021.05.018.

42. Pagani, L., Lawson, D.J., Jagoda, E., Morseburg, A., Eriksson, A., Mitt, M., Clemente, F., Hudjashov, G., DeGiorgio, M., Saag, L., et al. (2016). Genomic analyses inform on migration events during the peopling of Eurasia. Nature 538, 238–242. 10.1038/nature19792.

43. Wang, B., Zhang, Y.B., Zhang, F., Lin, H., Wang, X., Wan, N., Ye, Z., Weng, H., Zhang, L., Li, X., et al. (2011). On the origin of Tibetans and their genetic basis in adapting high-altitude environments. PLoS One 6, e17002, e17002. 10.1371/journal.pone.0017002.

44. Wuren, T., Simonson, T.S., Qin, G., Xing, J., Huff, C.D., Witherspoon, D.J., Jorde, L.B., and Ge, R.L. (2014). Shared and unique signals of high-altitude adaptation in geographically distinct Tibetan populations. PLoS One 9, e88252. 10.1371/journal.pone.0088252.

45. Xu, S., Li, S., Yang, Y., Tan, J., Lou, H., Jin, W., Yang, L., Pan, X., Wang, J., Shen, Y., et al. (2011). A genome-wide search for signals of high-altitude adaptation in Tibetans. Mol Biol Evol 28, 1003–1011. 10.1093/molbev/msq277.

46. Yi, X., Liang, Y., Huerta-Sanchez, E., Jin, X., Cuo, Z.X., Pool, J.E., Xu, X., Jiang, H., Vinckenbosch, N., Korneliussen, T.S., et al. (2010). Sequencing of 50 human exomes reveals adaptation to high altitude. Science 329, 75–78. 10.1126/science.1190371.

47. Ben-Eghan, C., Sun, R., Hleap, J.S., Diaz-Papkovich, A., Munter, H.M., Grant, A.V., Dupras, C., and Gravel, S. (2020). Don’t ignore genetic data from minority populations. Nature 585, 184–186. 10.1038/d41586-020-02547-3.

48. Bergstrom, A., McCarthy, S.A., Hui, R., Almarri, M.A., Ayub, Q., Danecek, P., Chen, Y., Felkel, S., Hallast, P., Kamm, J., et al. (2020). Insights into human genetic variation and population history from 929 diverse genomes. Science 367. 10.1126/science.aay5012.

49. Choin, J., Mendoza-Revilla, J., Arauna, L.R., Cuadros-Espinoza, S., Cassar, O., Larena, M., Ko, A.M., Harmant, C., Laurent, R., Verdu, P., et al. (2021). Genomic insights into population history and biological adaptation in Oceania. Nature 592, 583–589. 10.1038/s41586-021-03236-5.

50. Liu, Y., Xie, J., Wang, M., Liu, C., Zhu, J., Zou, X., Li, W., Wang, L., Leng, C., Xu, Q., et al. (2021). Genomic Insights Into the Population History and Biological Adaptation of Southwestern Chinese Hmong-Mien People. Front Genet 12, 815160. 10.3389/fgene.2021.815160.

51. Li, P. (2010). Recognition of History and Culture and Identification of Yi people. Journal of Guizhou University for National Titles (Philosophy and Social Science).

52. Liu, C.C., Witonsky, D., Gosling, A., Lee, J.H., Ringbauer, H., Hagan, R., Patel, N., Stahl, R., Novembre, J., Aldenderfer, M., et al. (2022). Ancient genomes from the Himalayas illuminate the genetic history of Tibetans and their Tibeto-Burman speaking neighbors. Nat Commun 13, 1203. 10.1038/s41467-022-28827-2.

53. Wang, C.C., Wang, L.X., Shrestha, R., Zhang, M., Huang, X.Y., Hu, K., Jin, L., and Li, H. (2014). Genetic structure of Qiangic populations residing in the western Sichuan corridor. PLoS One 9, e103772. 10.1371/journal.pone.0103772.

54. Kamberov, Y.G., Wang, S., Tan, J., Gerbault, P., Wark, A., Tan, L., Yang, Y., Li, S., Tang, K., Chen, H., et al. (2013). Modeling recent human evolution in mice by expression of a selected EDAR variant. Cell 152, 691–702. 10.1016/j.cell.2013.01.016.

55. Basu Mallick, C., Iliescu, F.M., Mols, M., Hill, S., Tamang, R., Chaubey, G., Goto, R., Ho, S.Y., Gallego Romero, I., Crivellaro, F., et al. (2013). The light skin allele of SLC24A5 in South Asians and Europeans shares identity by descent. PLoS Genet 9, e1003912. 10.1371/journal.pgen.1003912.

56. Zhang, X., Sun, A., and Ge, J. (2021). Origin and Spread of the ALDH2 Glu504Lys Allele. Phenomics 1, 222–228. 10.1007/s43657-021-00017-y.

57. Smith, J.D., Meehan, M.H., Crean, J., and McCann, A. (2011). Alpha T-catenin (CTNNA3): a gene in the hand is worth two in the nest. Cell Mol Life Sci 68, 2493–2498. 10.1007/s00018-011-0728-0.

58. Chouk, H., Ben Rejeb, M., Boussofara, L., Elmabrouk, H., Ghariani, N., Sriha, B., Saad, A., H’Mida, D., and Denguezli, M. (2021). Phenotypic intrafamilial variability including H syndrome and Rosai-Dorfman disease associated with the same c.1088G > A mutation in the SLC29A3 gene. Human genomics 15, 63. 10.1186/s40246-021-00362-z.

59. Clark, M.B., Wrzesinski, T., Garcia, A.B., Hall, N.A.L., Kleinman, J.E., Hyde, T., Weinberger, D.R., Harrison, P.J., Haerty, W., and Tunbridge, E.M. (2020). Long-read sequencing reveals the complex splicing profile of the psychiatric risk gene CACNA1C in human brain. Molecular psychiatry 25, 37–47. 10.1038/s41380-019-0583-1.

60. Xu, Q., Miao, Y., Ren, J., Sun, Y., Li, C., Cai, X., and Wang, Z. (2022). Silencing of Nesprin-2 inhibits the differentiation of myofibroblasts from fibroblasts induced by mechanical stretch. Int Wound J 19, 978–986. 10.1111/iwj.13694.

61. Ni, M., Chen, Y., Sun, X., Deng, Y., Wang, X., Zhang, T., Wu, Y., Yu, L., Xu, S., Yu, H., et al. (2022). DNA methylation and transcriptional profiles of IRF5 gene in ankylosing spondylitis: A case-control study. Int Immunopharmacol 110, 109033. 10.1016/j.intimp.2022.109033.

62. Sha, X., Wang, K., Wang, F., Zhang, C., Yang, L., and Zhu, X. (2021). Silencing PFKP restrains the stemness of hepatocellular carcinoma cells. Exp Cell Res 407, 112789. 10.1016/j.yexcr.2021.112789.

63. Opalinski, L., Veenhuis, M., and van der Klei, I.J. (2011). Peroxisomes: membrane events accompanying peroxisome proliferation. Int J Biochem Cell Biol 43, 847–851. 10.1016/j.biocel.2011.03.006.

64. Liao, W.W., Asri, M., Ebler, J., Doerr, D., Haukness, M., Hickey, G., Lu, S., Lucas, J.K., Monlong, J., Abel, H.J., et al. (2023). A draft human pangenome reference. Nature 617, 312–324. 10.1038/s41586-023-05896-x.

65. Gao, Y., Yang, X., Chen, H., Tan, X., Yang, Z., Deng, L., Wang, B., Kong, S., Li, S., Cui, Y., et al. (2023). A pangenome reference of 36 Chinese populations. Nature. 10.1038/s41586-023-06173-7.

66. 66. Zhu, K., Du, P., Li, J., Zhang, J., Hu, X., Meng, H., Chen, L., Zhou, B., Yang, X., Xiong, J., et al. (2022). Cultural and demic co-diffusion of Tubo Empire on Tibetan Plateau. iScience 25. 10.1016/j.isci.2022.105636.

67. Gao, Y., Zhang, X., Chen, H., Lu, Y., Ma, S., Yang, Y., Zhang, M., and Xu, S. (2022). Reconstructing the ancestral gene pool to uncover the origins and genetic links of Hmong–Mien speakers. Cell Genomics.

68. Pan, Y., Zhang, C., Lu, Y., Ning, Z., Lu, D., Gao, Y., Zhao, X., Yang, Y., Guan, Y., Mamatyusupu, D., and Xu, S. (2022). Genomic diversity and post-admixture adaptation in the Uyghurs. Natl Sci Rev 9, nwab124. 10.1093/nsr/nwab124.

69. LaPolla, R.J. (2013). 25 Eastern Asia: Sino - Tibetan linguistic history. In The Encyclopedia of Global Human Migration. 10.1002/9781444351071.wbeghm825.

70. Jeong, C. (2017). A longitudinal cline characterizes the genetic structure of human populations in the Tibetan plateau. PLoS ONE 12.

71. Jeong, C., Alkorta-Aranburu, G., Basnyat, B., Neupane, M., Witonsky, D.B., Pritchard, J.K., Beall, C.M., and Di Rienzo, A. (2014). Admixture facilitates genetic adaptations to high altitude in Tibet. Nat Commun 5, 3281. 10.1038/ncomms4281.

72. Tao, L. (2022). Ancient Genomes Reveal Coexistence of Demic and Cultural Diffusion in the Development of Neolithic Mixed Millet and Rice Farming in Southwest China. Current Biology.

73. Galanter, J.M., Gignoux, C.R., Oh, S.S., Torgerson, D., Pino-Yanes, M., Thakur, N., Eng, C., Hu, D., Huntsman, S., Farber, H.J., et al. (2017). Differential methylation between ethnic sub-groups reflects the effect of genetic ancestry and environmental exposures. eLife 6. 10.7554/eLife.20532.

74. Peng, Y., Cui, C., He, Y., Ouzhuluobu, Zhang, H., Yang, D., Zhang, Q., Bianbazhuoma, Yang, L., He, Y., et al. (2017). Down-Regulation of EPAS1 Transcription and Genetic Adaptation of Tibetans to High-Altitude Hypoxia. Mol Biol Evol 34, 818–830. 10.1093/molbev/msw280.

75. Yang, Z., Bai, C., Pu, Y., Kong, Q., Guo, Y., Ouzhuluobu, Gengdeng, Liu, X., Zhao, Q., Qiu, Z., et al. (2022). Genetic adaptation of skin pigmentation in highland Tibetans. Proc Natl Acad Sci U S A 119, e2200421119. 10.1073/pnas.2200421119.

76. Lin, Z., Lu, Y., Yu, G., Teng, H., Wang, B., Yang, Y., Li, Q., Sun, Z., Xu, S., Wang, W., and Tian, P. (2023). Genome-wide DNA methylation landscape of four Chinese populations and epigenetic variation linked to Tibetan high-altitude adaptation. Sci China Life Sci. 10.1007/s11427-022-2284-8.

77. World Medical Association, I. (2001). World Medical Association Declaration of Helsinki. Ethical principles for medical research involving human subjects. Bull World Health Organ 2001.

78. Chang, C.C., Chow, C.C., Tellier, L.C., Vattikuti, S., Purcell, S.M., and Lee, J.J. (2015). Second-generation PLINK: rising to the challenge of larger and richer datasets. Gigascience 4, 7. 10.1186/s13742-015-0047-8.

79. Manichaikul, A., Mychaleckyj, J.C., Rich, S.S., Daly, K., Sale, M., and Chen, W.M. (2010). Robust relationship inference in genome-wide association studies. Bioinformatics 26, 2867–2873. 10.1093/bioinformatics/btq559.

80. Almarri, M.A., Bergstrom, A., Prado-Martinez, J., Yang, F., Fu, B., Dunham, A.S., Chen, Y., Hurles, M.E., Tyler-Smith, C., and Xue, Y. (2020). Population Structure, Stratification, and Introgression of Human Structural Variation. Cell 182, 189–199 e115. 10.1016/j.cell.2020.05.024.

81. Byrska-Bishop, M., Evani, U.S., Zhao, X., Basile, A.O., Abel, H.J., Regier, A.A., Corvelo, A., Clarke, W.E., Musunuri, R., Nagulapalli, K., et al. (2022). High-coverage whole-genome sequencing of the expanded 1000 Genomes Project cohort including 602 trios. Cell 185, 3426–3440 e3419. 10.1016/j.cell.2022.08.004.

82. Patterson, N., Moorjani, P., Luo, Y., Mallick, S., Rohland, N., Zhan, Y., Genschoreck, T., Webster, T., and Reich, D. (2012). Ancient admixture in human history. Genetics 192, 1065–1093. 10.1534/genetics.112.145037.

83. Bergström, A., McCarthy, S.A., Hui, R., Almarri, M.A., Ayub, Q., Danecek, P., Chen, Y., Felkel, S., Hallast, P., Kamm, J., et al. (2019). Insights into human genetic variation and population history from 929 diverse genomes. 10.1101/674986.

84. Patterson, N., Price, A.L., and Reich, D. (2006). Population structure and eigenanalysis. PLoS Genet 2, e190, e190. 10.1371/journal.pgen.0020190.

85. Alexander, D.H., Novembre, J., and Lange, K. (2009). Fast model-based estimation of ancestry in unrelated individuals. Genome Res 19, 1655–1664. 10.1101/gr.094052.109.

86. Reich, D., Thangaraj, K., Patterson, N., Price, A.L., and Singh, L. (2009). Reconstructing Indian population history. Nature 461, 489–494. 10.1038/nature08365.

87. Delaneau, O., Marchini, J., and Zagury, J.F. (2011). A linear complexity phasing method for thousands of genomes. Nat Methods 9, 179–181. 10.1038/nmeth.1785.

88. Browning, S.R., and Browning, B.L. (2015). Accurate Non-parametric Estimation of Recent Effective Population Size from Segments of Identity by Descent. Am J Hum Genet 97, 404–418. 10.1016/j.ajhg.2015.07.012.

89. Browning, B.L., and Browning, S.R. (2013). Detecting identity by descent and estimating genotype error rates in sequence data. Am J Hum Genet 93, 840–851. 10.1016/j.ajhg.2013.09.014.

90. Pickrell, J.K., and Pritchard, J.K. (2012). Inference of population splits and mixtures from genome-wide allele frequency data. PLoS Genet 8, e1002967. 10.1371/journal.pgen.1002967.

91. Loh, P.R., Lipson, M., Patterson, N., Moorjani, P., Pickrell, J.K., Reich, D., and Berger, B. (2013). Inferring admixture histories of human populations using linkage disequilibrium. Genetics 193, 1233–1254. 10.1534/genetics.112.147330.

92. Lawson, D.J., Hellenthal, G., Myers, S., and Falush, D. (2012). Inference of population structure using dense haplotype data. PLoS Genet 8, e1002453. 10.1371/journal.pgen.1002453.

