## supplemental files.pdf for "Distinguished biological adaptation architecture aggravated population differentiation of Tibeto-Burman-speaking people inferred from 500 whole-genome data from 39 populations": supplemental files.pdf

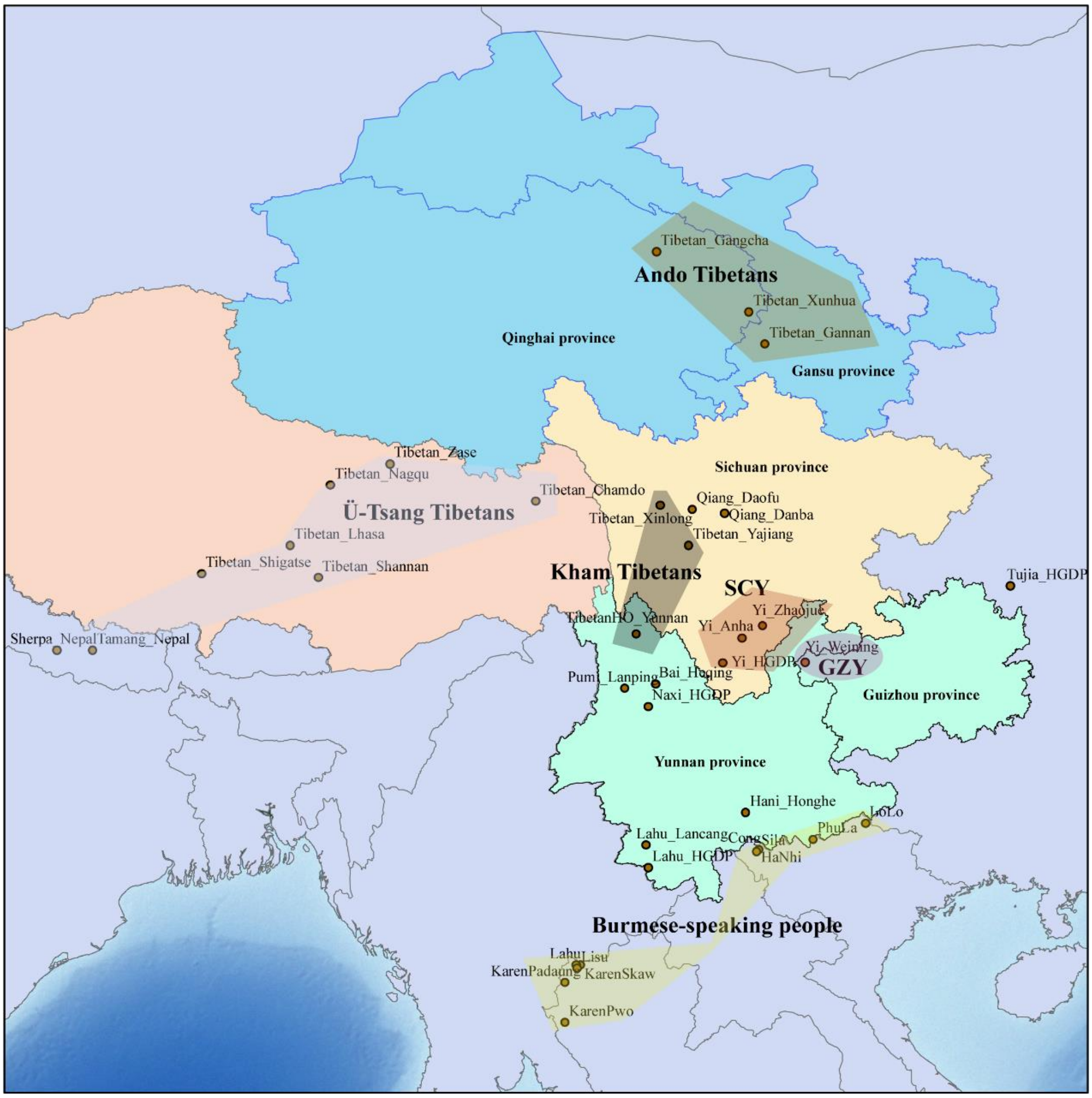


**Figure S1. Sample location of TB populations**

Sampling map based on 500 individuals from 39 TB populations. The distribution is mainly in southwestern China and northern mainland Southeast Asia, including the Tibetan plateau (TP), the Tibetan-Yi corridor (TYC), the Yunnan-Guizhou plateau (YGP), the Alluvial plains (AP) and Deltas, etc. geographical patterns.

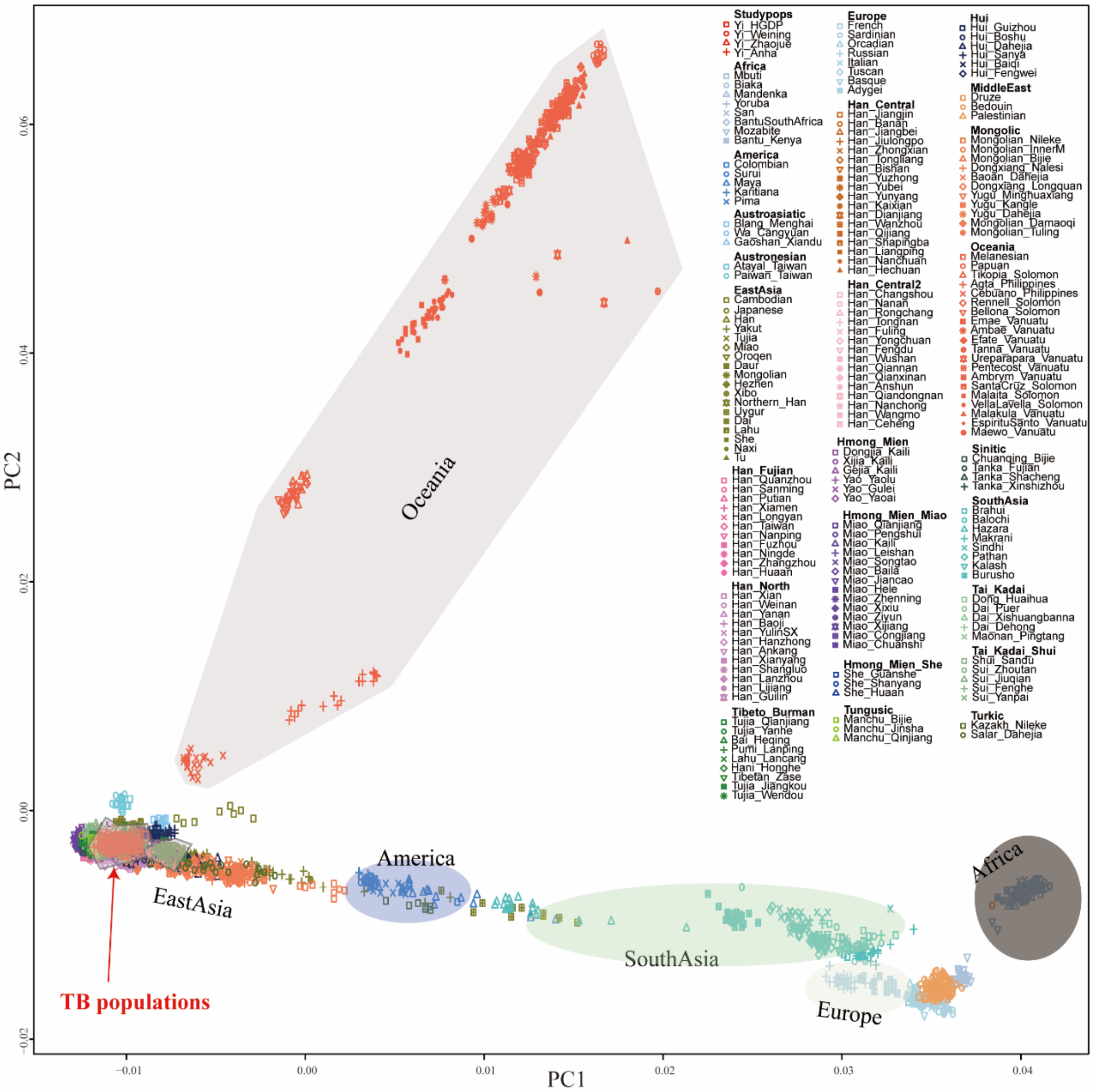


**Figure S2. The genetic location of TB populations at the world-scale by Principal component analysis (PCA)**

Principal component analysis (PCA) at the world-scale based on high-density datasets, with TB populations and East Asian populations clustering with each other.

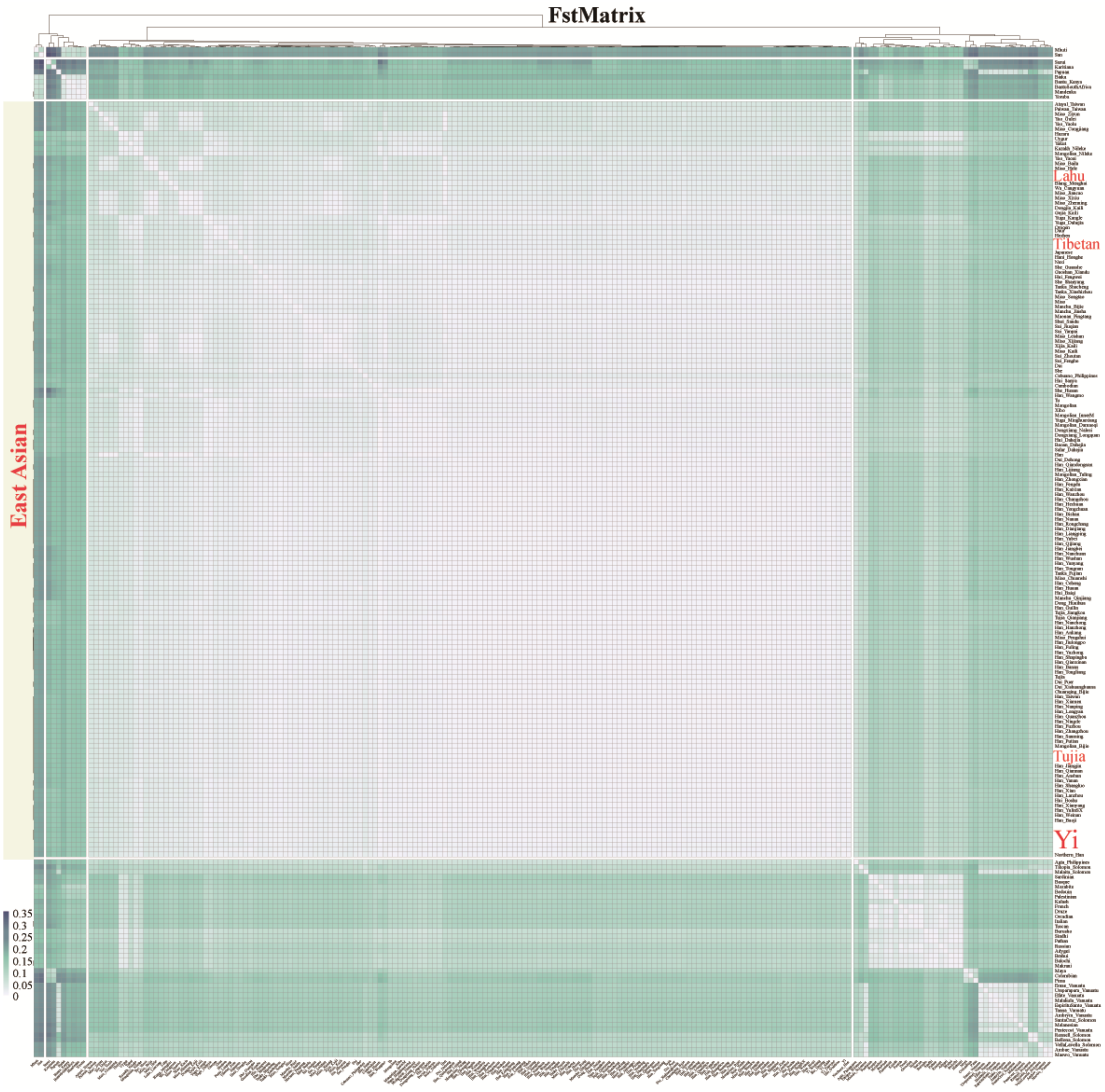


**Figure S3. The genetic affinity of TB populations at the world-scale by Fst Matrix**

The Fst Matrix at the world-scale, with East Asian populations in the middle, shows smaller Fst values, indicating closer genetic affinity.

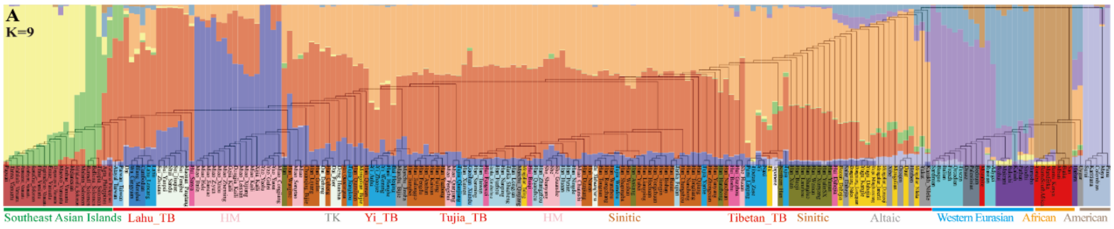


**Figure S4. Phylogenetic structure of TB populations at world-scale**

Composition graph of Neighbor-joining Tree generated by Fst and ADMIXTURE based on the high-density dataset.

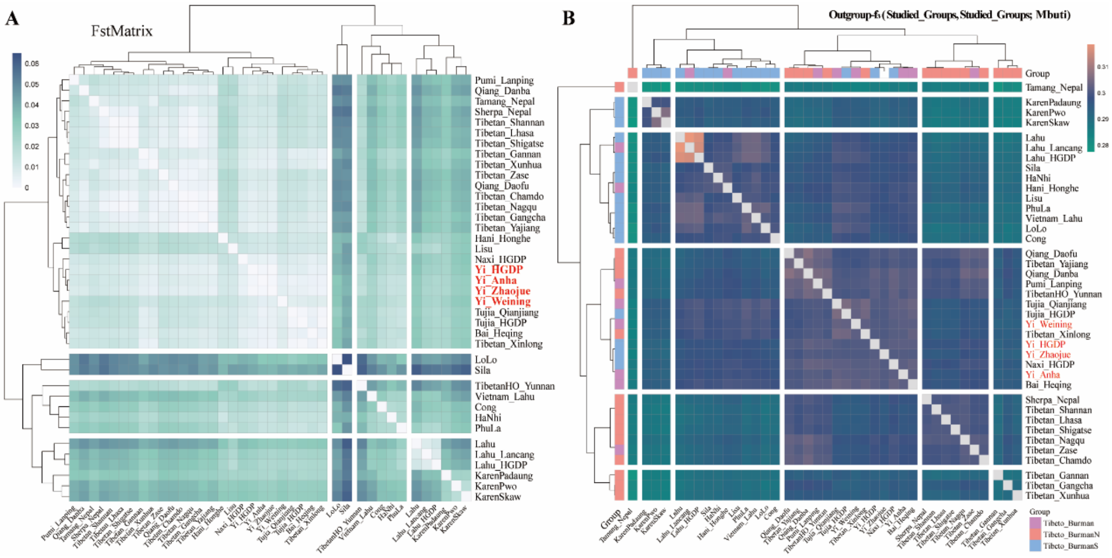


**Figure S5. Genetic affinity of TB populations at the TB scale**

Genetic affinity of TB populations observed based on low-density datasets. (A) Fst Matrix heat map from top to bottom for TPT, ethnics from TYC and YGP and APB populations, respectively. (B) Outgroup-$f_3$ heatmap (TB population, TB population; Mbuti), with higher Outgroup-$f_3$ values indicating closer relatedness.

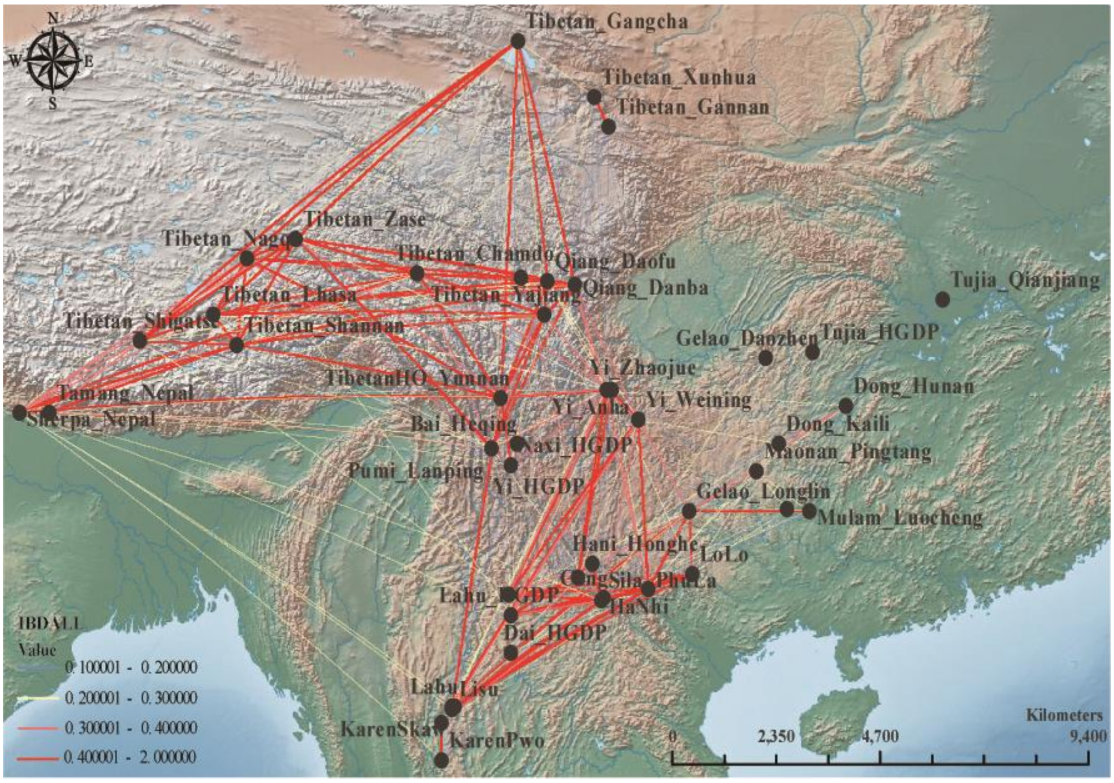


**Figure S6. Identity-by-descent (IBD) map represents the interactive pattern of TB populations**
Visualization chart based on IBD all values to assess affinity among TB peoples. Affinity is represented by red, orange, yellow and blue from highest to lowest affinity.

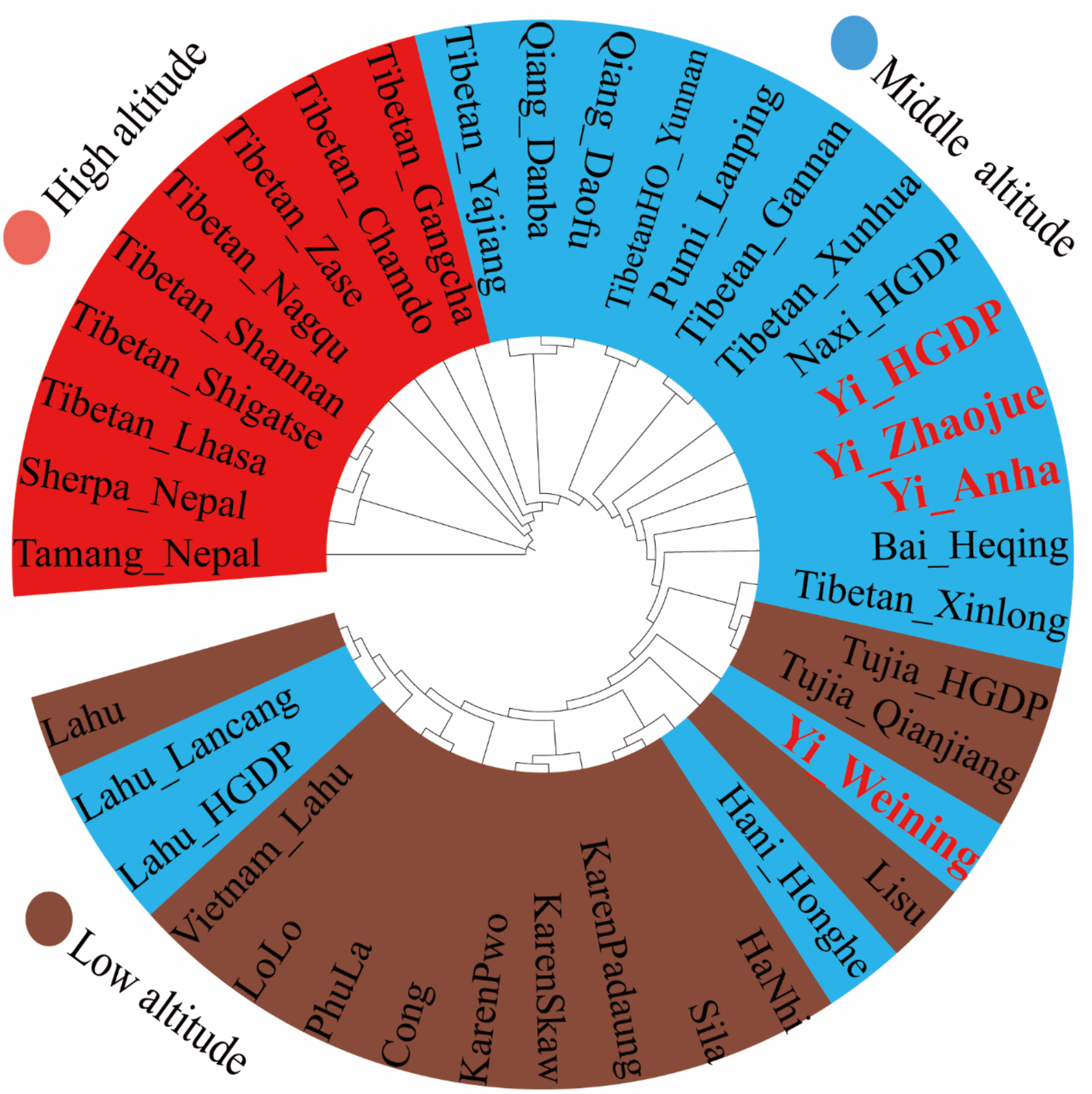


**Figure S7. Phylogenetic relationship generated by 1-Outgroup-$f_3$**

The Phylogenetic of 1-Outgroup-$f_3$ shows distribution according to high, middle and low altitude geographical patterns.

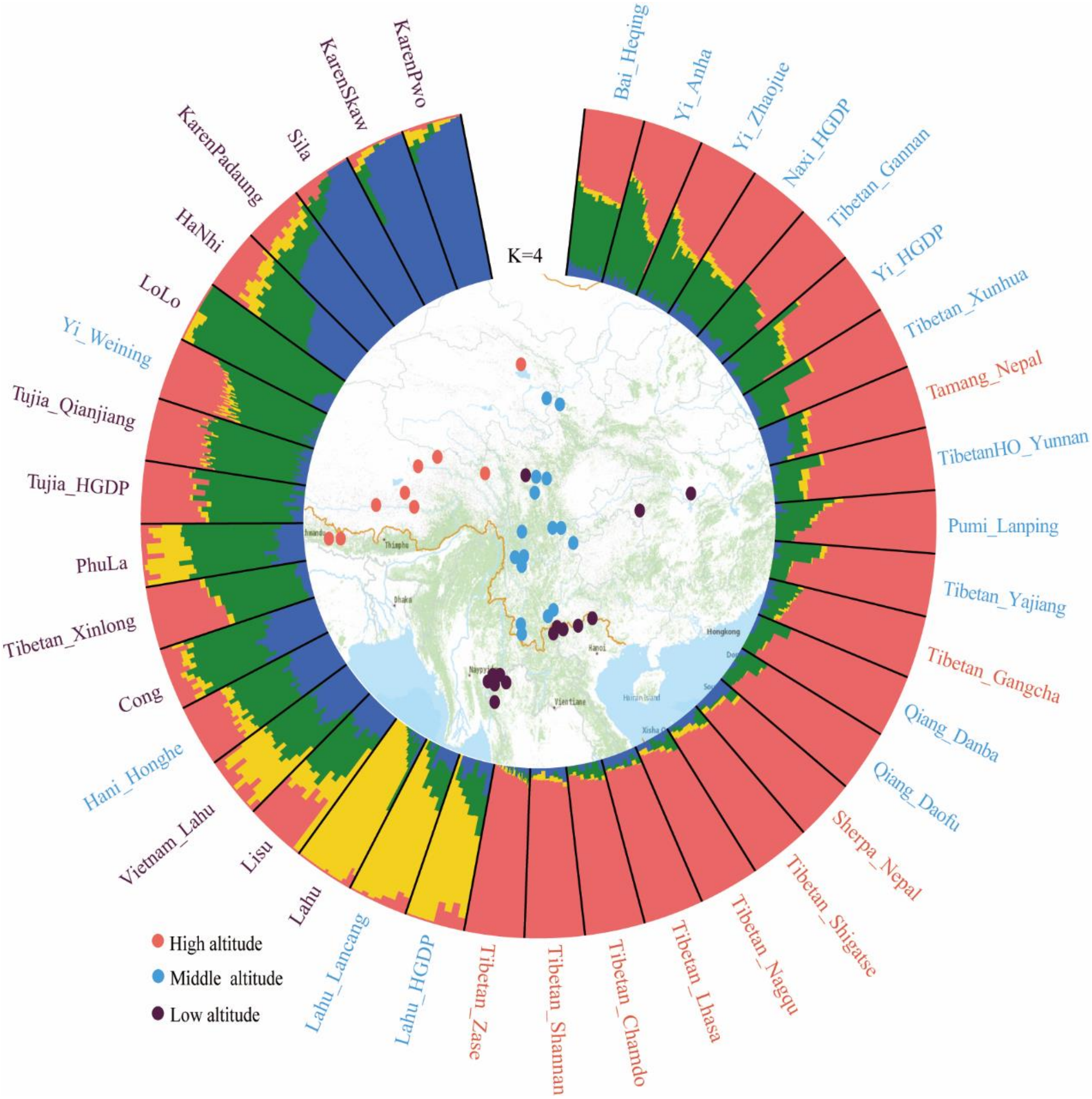


**Figure S8. The relevance between genetic affinity and geographical pattern.**
The outer circle shows the results of ADMIXTURE for TB populations based on K=4, and the inner circle shows a sampling map from 39 TB populations, differentiated by altitude markers.

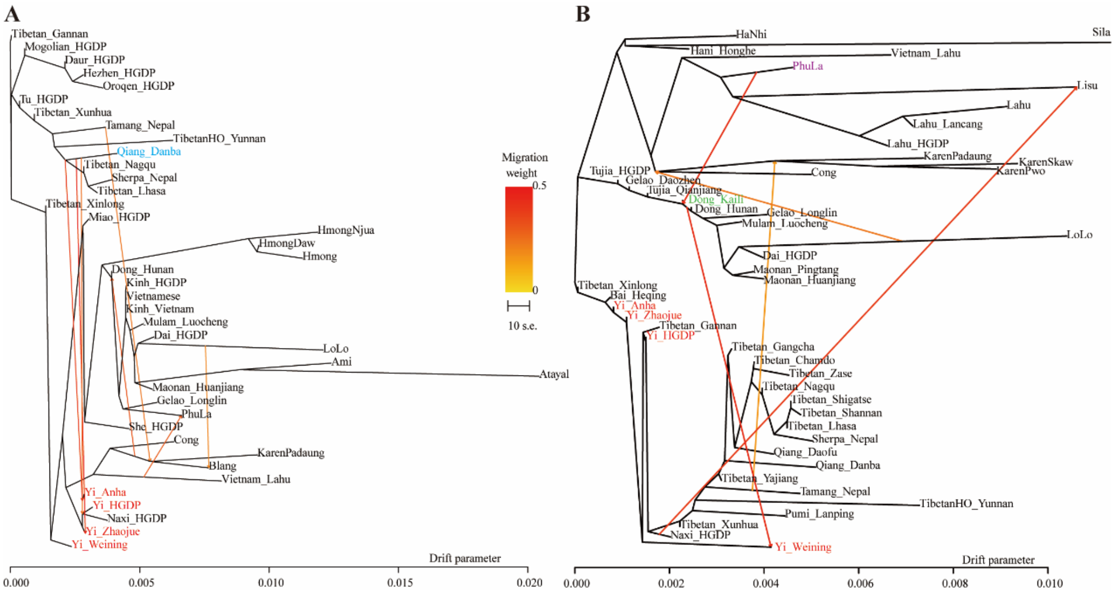


**Figure S9. Gene flow event observed by TreeMix**

A line with the arrow represents gene flow events from reference populations to studied populations.

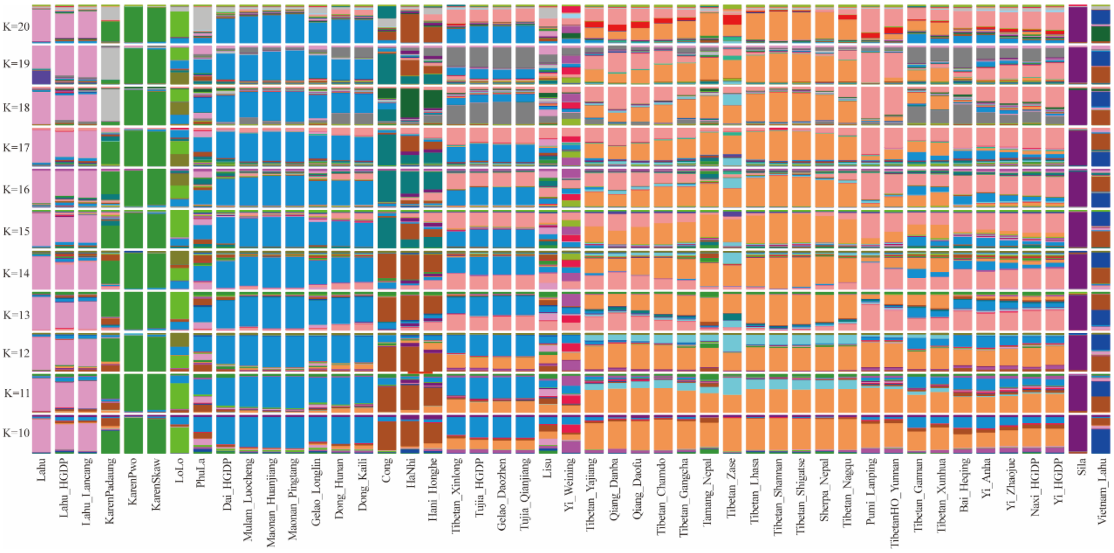


**Figure S10. Mean results of ADMIXTURE with K from 10 to 20**

We compared the mean results of ADMIXTURE with SCY (Yi_Zhaojue, Yi_Anha and Yi_HGDP) and GZY (Yi_Weining), finding that as the ancestral component grew, the proportion of GZY became increasingly different from SCY.

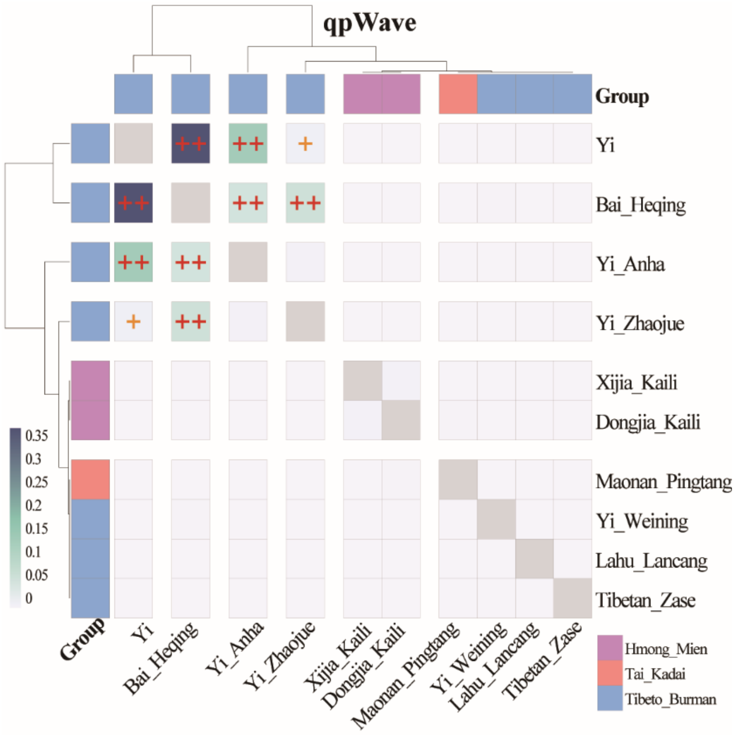


**Figure S11. Testing genetic homogeneity by qpWave**

The results demonstrate that SCY is homogeneous with Bai and heterogeneous with GZY. +: 0.05> P> 0.01, ++: P> 0.05

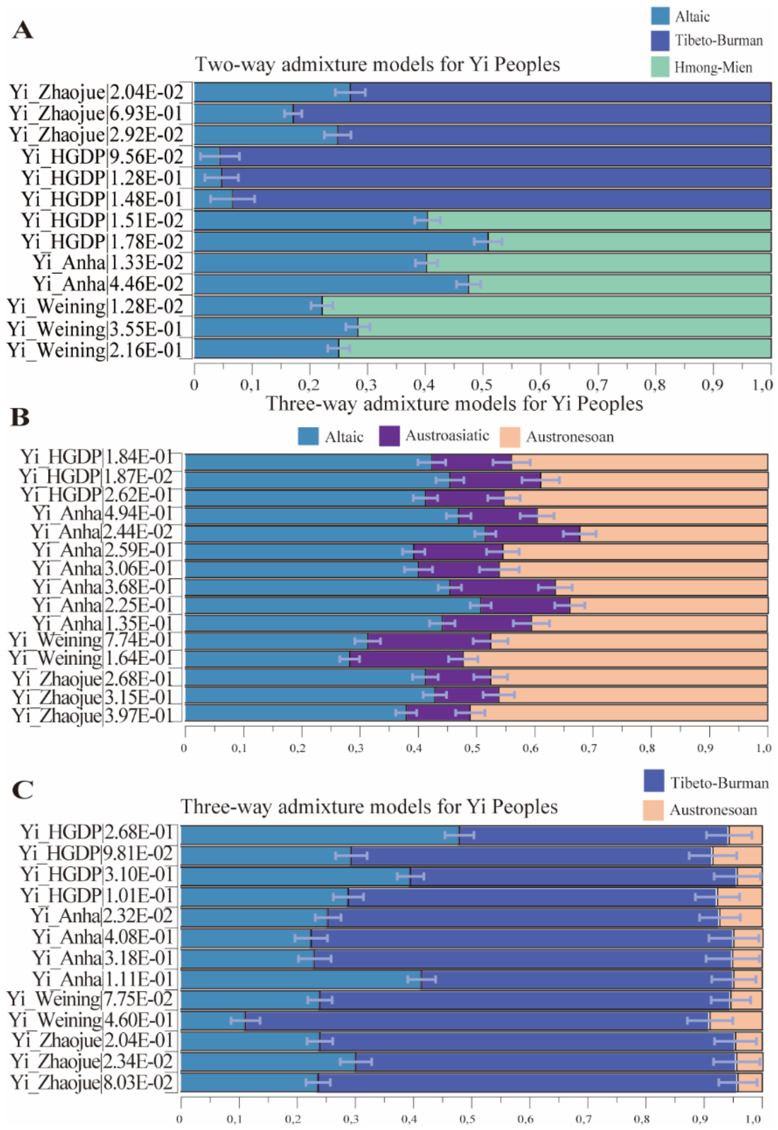


**Figure S12. Presenting the admixture model of Yi via qpAdm**

(A) Two-way admixture models for Yi Peoples. Light blue for the Altaic populations, navy for the Tibeto-Burman populations, and green for the Hmong-Mien populations. Outgroup: Cambodian. (B) and (C) Three-way admixture models for Yi Peoples. Purple for the Austroasiatic populations. Brick red for the Austronesian population. Outgroup: Cambodian. For detailed results, refer to Table S3-5.

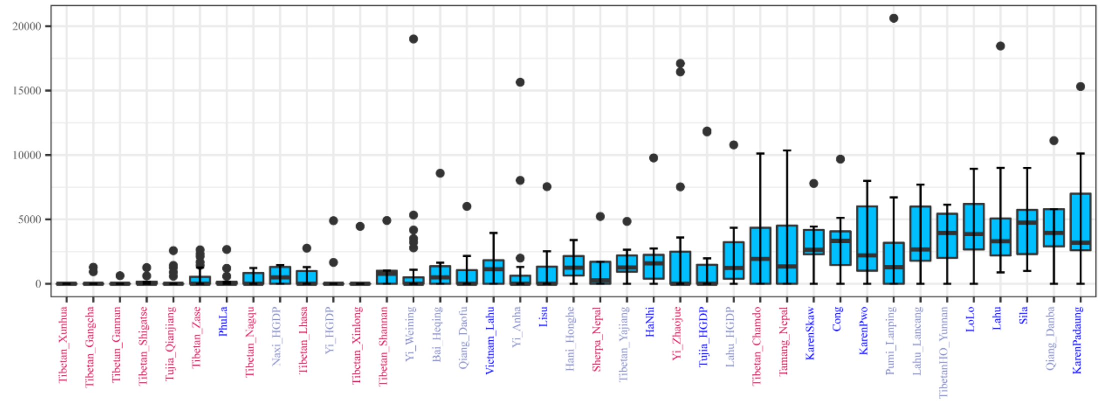


**Figure S13. Runs of homozygosity (ROH) of TB populations.**
The y-axis represents the number of ROH and the x-axis represents the populations. From left to right, the TPT, TYT and Ando Tibetans, Yi, Tujia and APB populations are shown from lowest to highest.

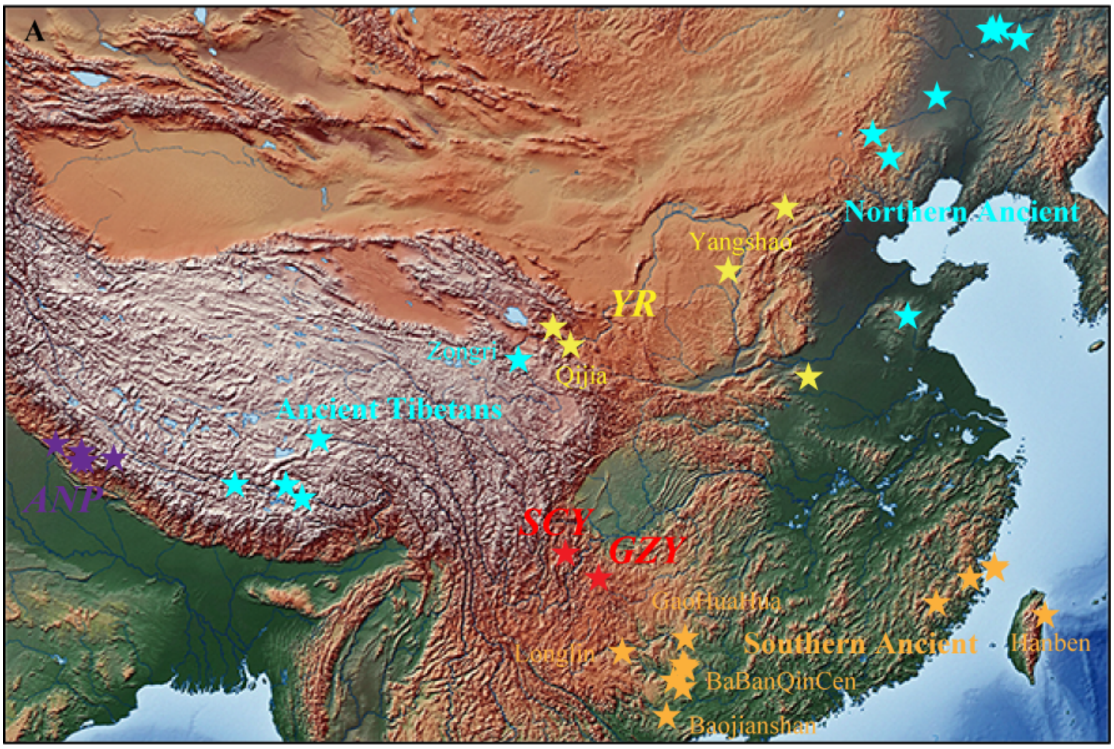


**Figure S14. Map of paleogenomics distribution of populations**
The Paleogene group is roughly divided into three categories. Northern ancients: including YR and Amur River populations. Southern ancients: including Fujian and Guangxi ancient populations. TP ancients: including Nepal and Plateau ancient populations.

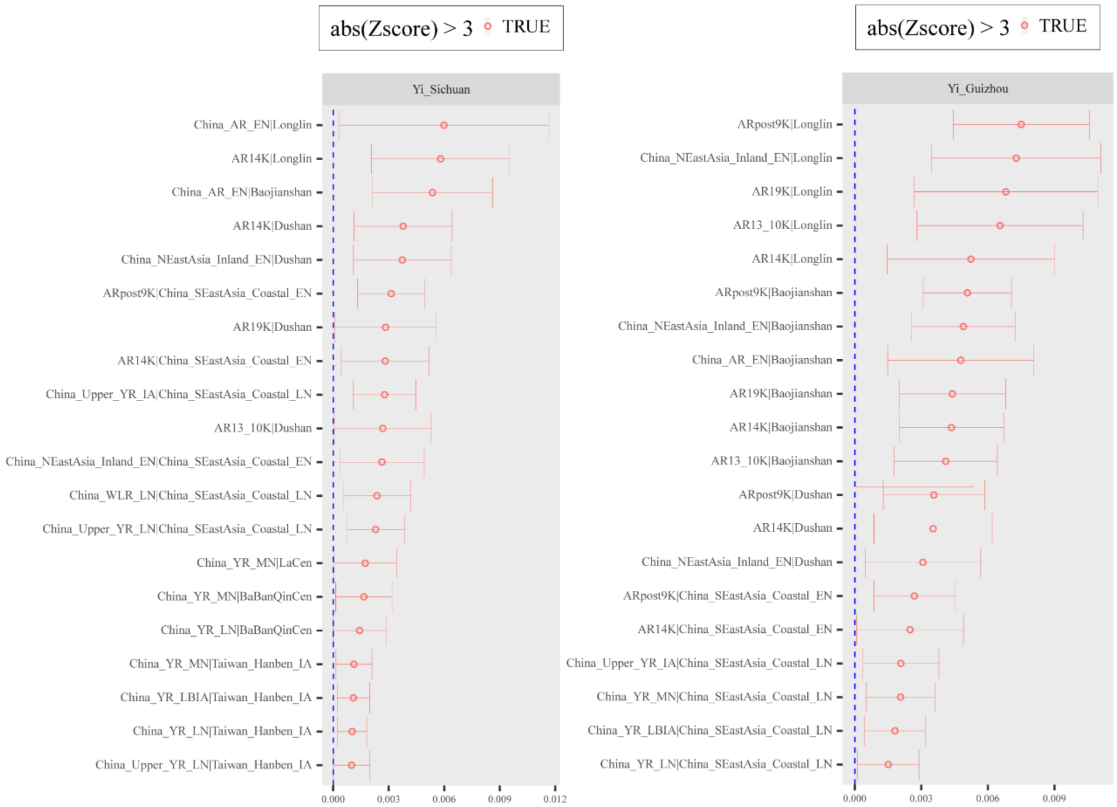


**Figure S15. Affinity testing of ancient populations.**

The genetic affinity between Yi and ancient populations measured by affinity $f_4$-statistic, which forms of (Northern Ancients, Southern Ancients; SCY/GZY, Mbuti). As shown in the lable the results are positive, with the larger results indicating that SCY/GZY enjoys more alleles and is closer in affinity to Northern Ancients than Southern Ancients.

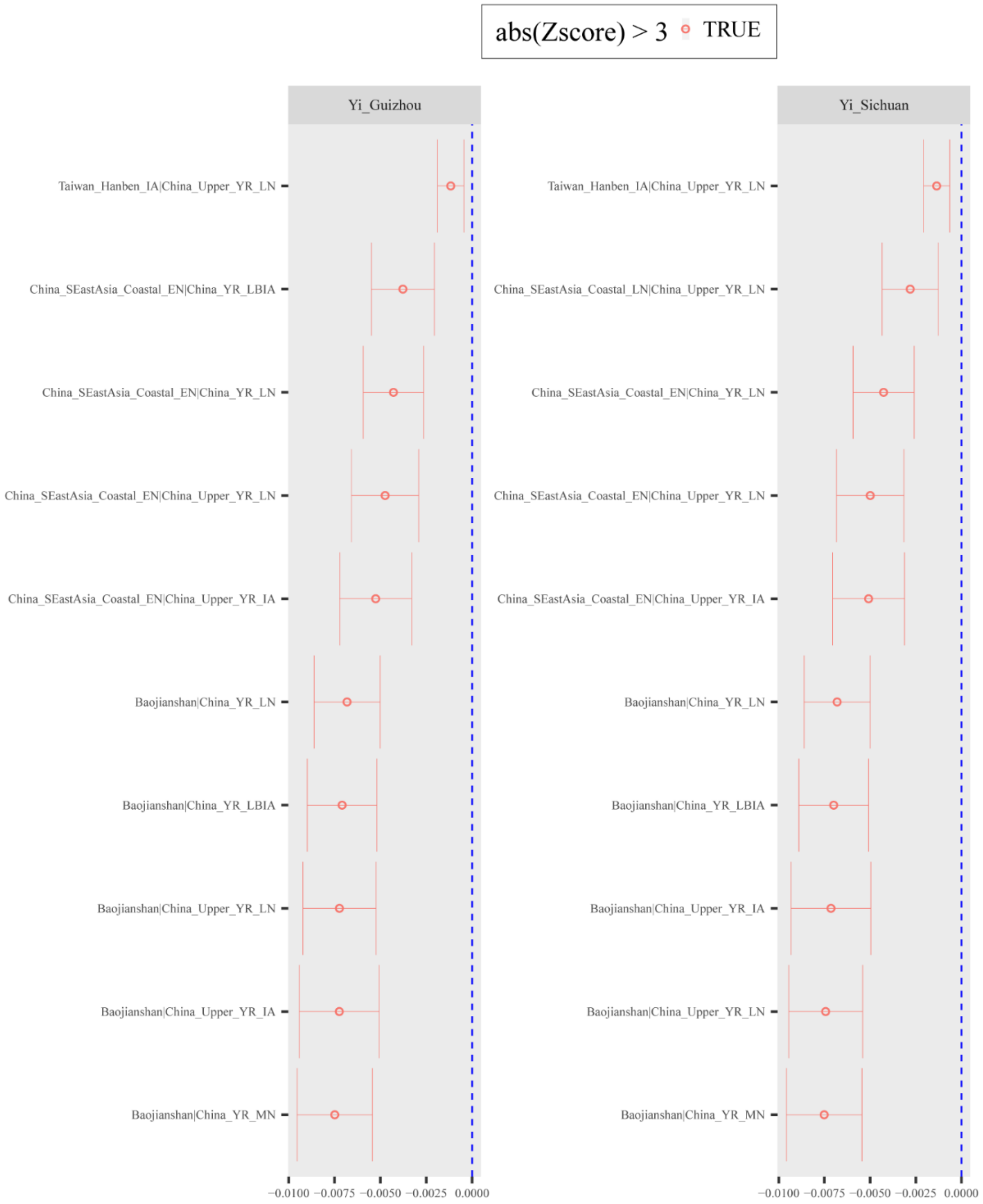


**Figure S16. Asymmetric $f_4$ tests to identify the Northern ancestral source**

Asymmetric $f_4$ tests were performed with forming (Southern Ancients, SCY/GZY; Northern Ancients, Mbuti), which showed significant negative scores, identifying the ancient populations of the Northern YR are the major ancestral source for Yi populations.

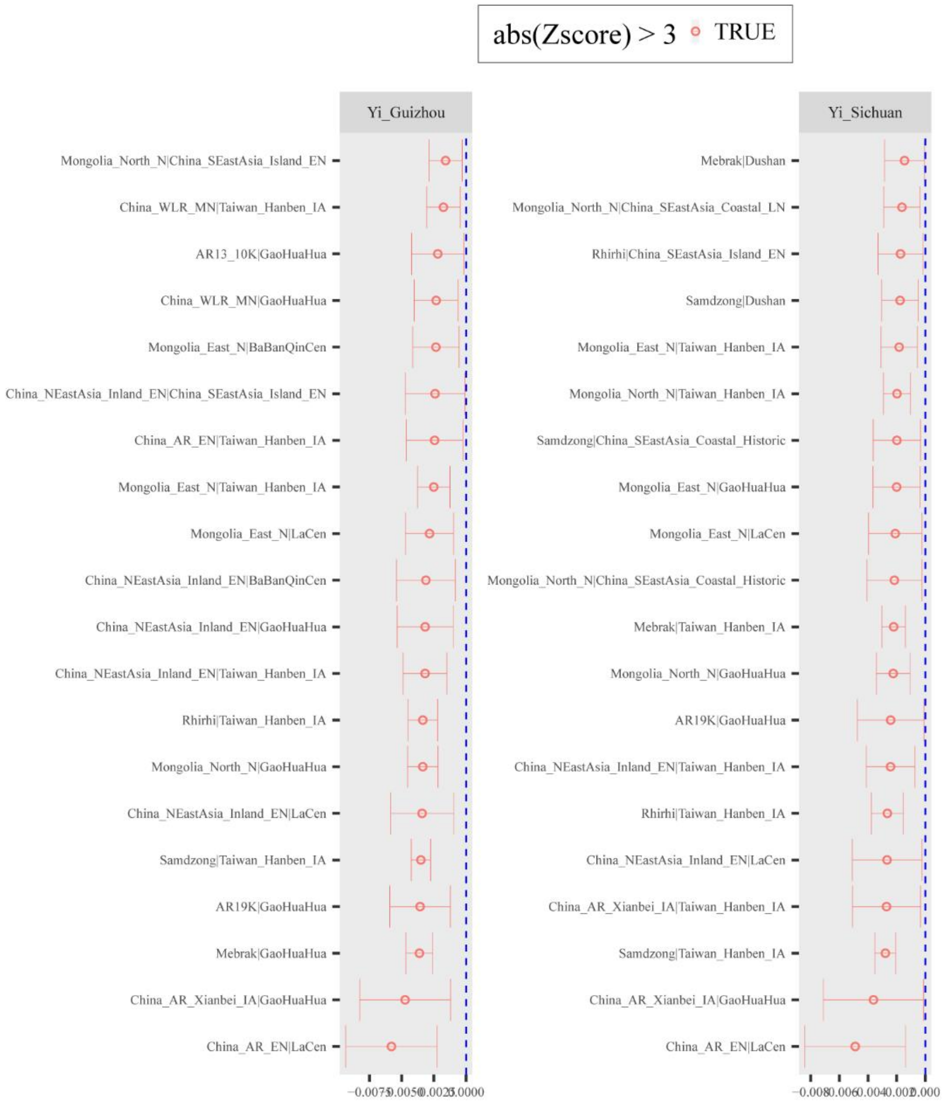


**Figure S17. Asymmetric *f*4 tests to identify the southern ancestral source**

Asymmetric *f*4 tests were generated with forming (Northern Ancients besides YR, SCY/GZY; Southern Ancients, Mbuti), which display significant negative values, demonstrating that southern ancients influenced the Yi populations.

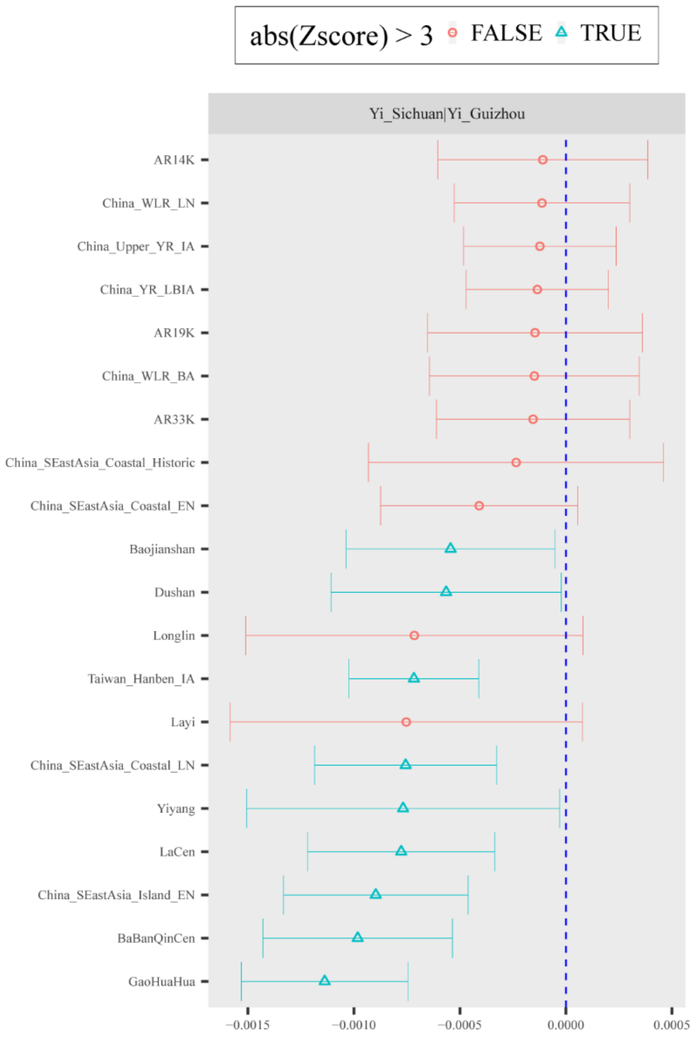


**Figure S18. Symmetry $f_4$ analysis revealing GZY was affected by ancient interactions located in southern China**

Symmetry $f_4$ analysis were generated with forming (SCY, GZY; Reference Ancients, Mbuti). As shown in the lable, more negative values of significance (green triangles) reveal that GZY was affected by ancient interactions in southern China frequently compared to SCY.

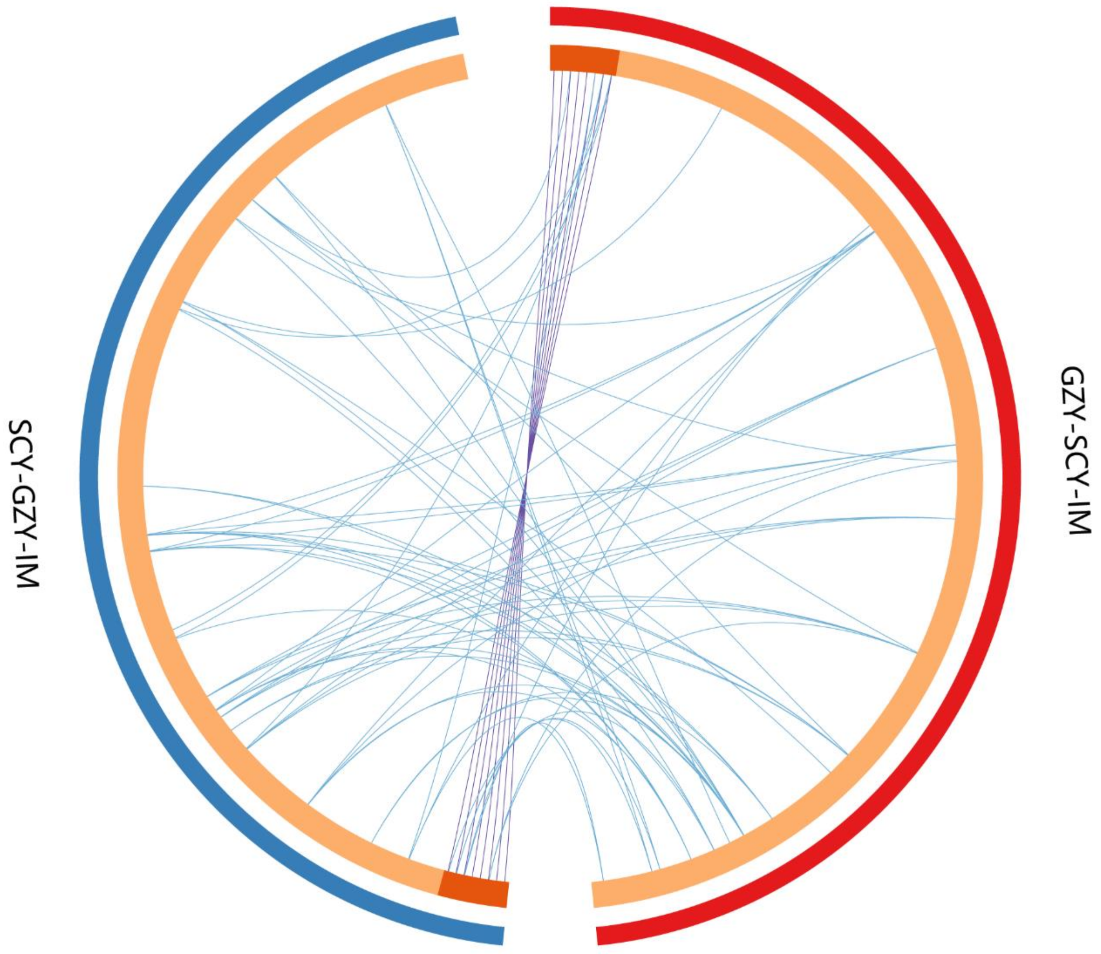


**Figure S19. Overlap between gene lists of SCY and GZY under local adaption**
At the gene level, purple curves connect genes that are identical and blue curves connect genes enriched for the same pathway. The inner circle represents the list of genes, with shared genes in orange and specific genes in yellow.

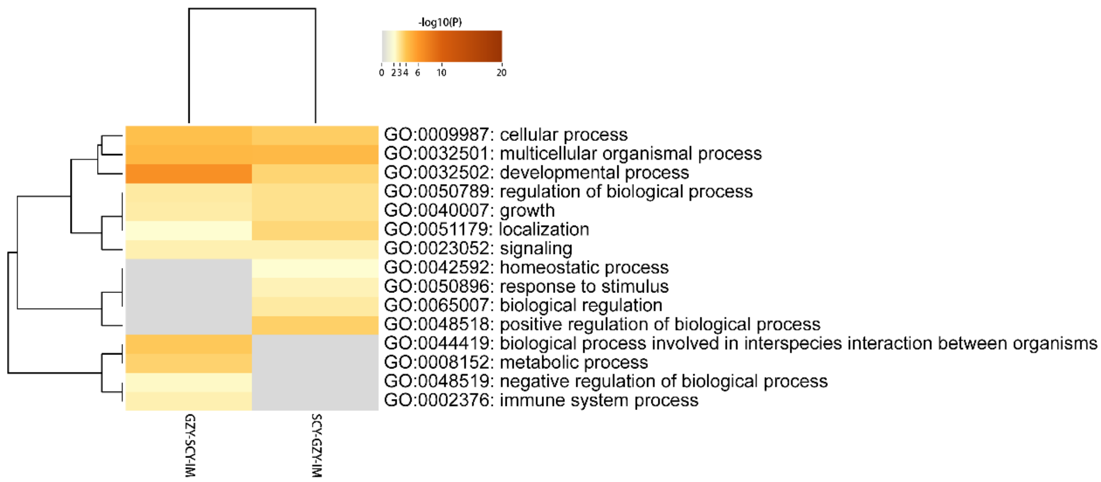


**Figure S20. The top-level Gene Ontology biological processes between SCY and GZY**

The branch on the left represents the clustering of enriched pathways. $-\log_{10}(P)$ values indicate how many genes the pathway is enriched by, with larger values (red) indicating more enriched genes and smaller values (white) indicating fewer enriched genes.
